# Probing interoception via thermosensation: No specific relationships across multiple interoceptive sub-modalities

**DOI:** 10.1101/2021.03.04.433866

**Authors:** Laura Crucianelli, Adam Enmalm, H. Henrik Ehrsson

**Affiliations:** Department of Neuroscience, Karolinska Institutet, Stockholm, Sweden

**Keywords:** temperature, homeostasis, body awareness, CT system, interoceptive battery

## Abstract

Interoception includes signals originating both from inside the body and from its surface, the skin. Here, we focused on the perception of temperature, a crucial modality for the maintenance of homeostasis. We used a classic (static) thermal detection task and developed a new dynamic *thermal matching task*, in which participants have to match a previously perceived moving thermal stimulus on the skin to a range of colder or warmer stimuli, presented in increasing or decreasing order. We investigated both hairy (forearm) and non-hairy (palm) skin, in keeping with previous tactile studies targeting the potential involvement of C-tactile fibres, which are part of an afferent homeostatic system found mainly on hairy skin. We also aimed at investigating the relationship between performance on the two thermal tasks and on three other tasks in different interoceptive sub-modalities: cardiac perception, affective touch, and pain detection. We found a significantly more accurate perception of dynamic temperature on hairy compared to non-hairy skin overall, particularly when the temperature was decreasing. Static perception of cooling was also superior on hairy skin and was related to dynamic temperature and pain only on non-hairy skin. Thus, our results suggest that hairy skin may have higher thermosensitivity than non-hairy skin and that dynamic thermosensation might offer a promising avenue to investigate thermosensation as a skin-based interoceptive submodality. Critically, we did not find any other significant relationship in performance among the four interoceptive modalities examined, which indicates independent processing and that interoception might be best quantified using a battery of tests.

## Introduction

Interoception has been defined as the body-to-brain axis of sensations concerning the state of the visceral body (Cameron, 2001; Sherrington, 1948), thus involving signals originating from within the body (e.g., cardiac, respiratory and digestive functions). However, physiological and anatomical observations led to a redefined and extended concept of interoception that encompasses information about the physiological condition of the entire body, including also signals originating from the body surface (e.g., temperature, itch, pain, and pleasure from sensual touch) and conveyed by specialised afferent pathways (Craig, 2002; Khalsa, Rudrauf, Feinstein, & Tranel, 2009). Interoception is related to the generation of bodily (affective) feelings, informing the organism about its bodily needs (Craig, 2008, 2009; Seth, 2013). Traditionally, interoception has been quantified using heartbeat detection tasks (Schandry, 1981), in which participants are asked to focus on their own heartbeats without touching their body but just by feeling the sensation of their heart beating. However, this approach has proved to be problematic (see Zamairola et al., 2018; Ainley et al., 2019; Zimpirich et al., 2019; Corneille et al., 2020 for a recent debate). Issues regarding this approach include evidence that performance in heartbeat detection tasks seems to be influenced by other factors, such as prior knowledge, heart rates, beliefs, practice, and even experimental instructions (e.g., Ring & Brener, 1996; Ring et al., 2015; Zamariola et al., 2018; Corneille et al., 2020). From a physiological point of view, the heart signal produces other bodily signals, such as vascular reactivity and muscle contractions, which can influence its perception. Thus, participants can quickly learn to use bodily strategies to alter, control, or manipulate the outcomes of heartbeat detection tasks (e.g., changes in respiration, tensing muscles, elicit stimuli to change heart rate, etc.; Ross & Brener, 1981; Whitehead & Drescher, 1980).

It has long been suggested that an optimal measure of interoception should be derived from visceral responses that are not highly correlated with skeletal muscle responses and thus potentially more difficult to be controlled by participants (Whitehead & Drescher, 1980). Indeed, interoception comprises other modalities that originate both inside the body and on the skin and provide information about the physiological status of the body at any given time (Craig, 2002; 2009). Thus, novel and perhaps more reliable methods to quantify interoception could potentially be developed above and beyond cardiac awareness.

In the last few decades, there has been an increasing focus on skin-mediated interoceptive modalities, such as pain and affective touch perception (Craig, 2002; 2003; Werner et al., 2009; Björnsdotter; Morrison and Olausson, 2010; Weiss et al., 2014; von Mohr & Fotopoulou, 2018). This interest has been partially motivated by the discovery of a specialised group of skin afferents, C Tactile afferents (CT; Vallbo, Olausson & Wessberg, 1999 for evidence in humans), that have been found in significantly larger amounts in the hairy skin of the body (Watkins, Dione, Ackerley, Backlund Wasling, Wessberg, & Löken, 2020). CT afferents have been proposed as the supporting system for the detection of affective touch (Löken et al, 2009; Morrison, Löken & Olausson, 2010), and they seem to project to the posterior insular cortex rather than to the somatosensory cortex (Björnsdotter, Löken, Olausson, Vallbo, & Wessberg, 2009 but see Gazzola et al. 2012). Importantly, the insula has been proposed to be a crucial area for interoceptive awareness and the experience of emotions (Critchley et al., 2004).

Although pain and affective touch are typically mediated by external causes and events occurring on the skin (even though painful and pleasant sensations can also originate from the skeletomuscular system, vascular system and inner organs), these modalities are homeostatically relevant since they provide information about physiological safety or threats (Craig, 2003; von Mohr & Fotopoulou, 2018). However, to our knowledge, less attention has been given to the perception of temperature as another skin-mediated interoceptive modality of fundamental importance for our survival (Craig et al., 2000; Craig, 2018, although thermal sensation can also arise from inside the body, as in the case of fever, Tabaren, 2018 for a review). The perception of peripheral temperature (or thermosensation) should be considered as distinct from the perception of the core temperature of the body, although these two aspects usually correlate and can influence each other and both contribute to the maintenance of thermoneutrality (Cabanac, Massonnet and Belaiche, 1972; Filingeri, Morris and Jay, 2017).

The perception of temperature is mediated by thermoreceptors, which are free nerve endings that signal the sensation of warmth and coolness (Sinclair, 1981; Abraira and Ginty, 2013; Filingeri, 2011; Jänig, 2018 for reviews). The skin is innervated by different types of afferent fibres encoding temperatures that can range from noxious cold to noxious heat, with innocuous cold and innocuous warm perception between the two extremes (Jänig, 2018 for a review). In particular, the perception of cold is mainly mediated via Aδ fibres (range ≈5-40°C; maximally discharging at approximately 30°C) and C fibres (e.g., Hensel et al., 1960; Iriuchijima and Zotterman, 1960; Iggo, 1969; Hensel and Wurster, 1969; Hensel and Iggo,1971; Darian-Smith et al., 1973; Dubner et al., 1975; Kenshalo and Duclaux, 1977; Jänig, 2018 for a review). Cooling (but not warming) of the skin also activates unmyelinated, low-threshold mechanoreceptors (CTs, Nordin, 1990). In contrast, warmth perception is mainly mediated by C fibres (range ≈29 - 45°C; maximally discharging at approximately 45°C, e.g., Iriuchijima and Zotterman, 1960, Hensel and Huopaniemi, 1969, Konietzny and Hensel, 1975; 1977; La Motte and Campbell, 1978; Hallin et al.; 1982). These C fibres also contribute to the sensations of pleasure and pain. Temperatures ≈< 15 and > 45°C activate cold and hot nociceptors, respectively (Kandel et al., 2000, Jänig, 2018; Table 1). Furthermore, there is an overlap between thermoreception and nociception, not only at the skin level but also in the spinal afferent pathway (Hellon, 1981, Marshall et al., 2019). Thus, given the challenges in disentangling the activation of mechanoreceptors, thermoreceptors and nociceptors, particularly in the interoceptive domain, only a few studies to date have explored how these stimuli interact in humans (e.g., Ackerley et al., 2018).

**Table 1.**
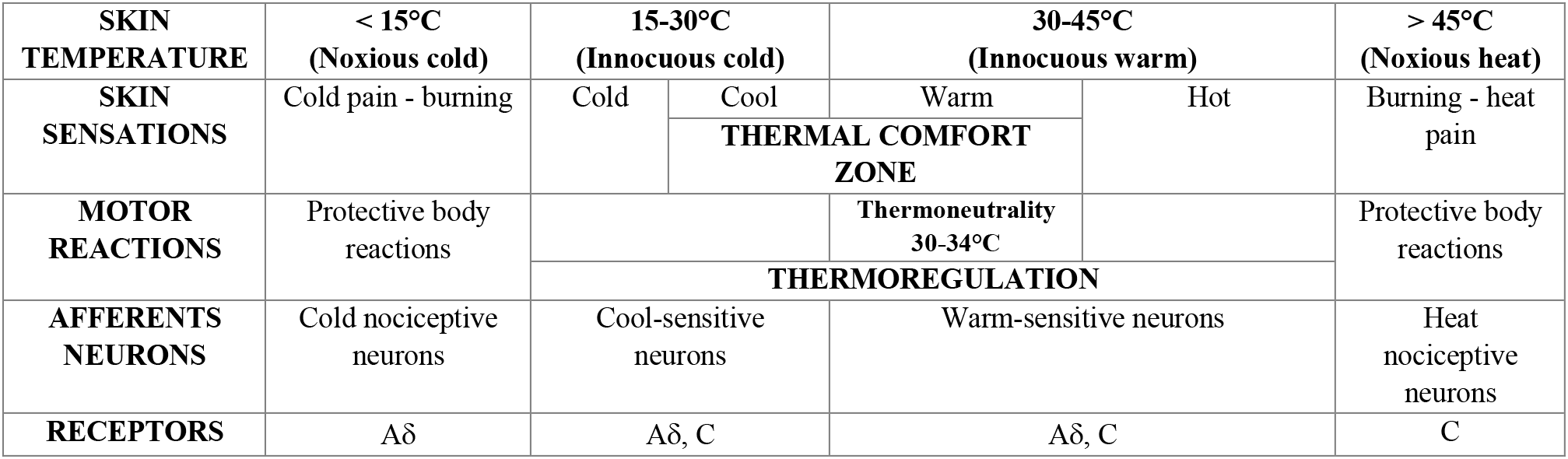
Activation of the different skin receptors in response to thermal stimuli. Adapted from Jänig, 2018

We normally conceptualise the perception of temperature as an exteroceptive modality, and it has long been considered as an aspect of touch (Sherrington, 1948). However, based on converging anatomical and physiological evidence thermosensation has been re-defined as interoceptive modality because it provides critical information about the body’s physiological state (Craig, 2003, 2008), and given its affective, functional and anatomical charactersitics (Cabanac et al., 1972; Mower, 1976; Craig, 2002). Here, we suggest that ad-hoc experimental studies that allow to keep the exteroceptive component of thermal stimulation (i.e., pressure and velocity of tactile stimulation) relatively constant, can target the interoceptive facet of this modality. In keeping with the aforementioned evidence supporting the interoceptive nature of some aspects of thermosensation (e.g., Craig, Chen, Bandy & Reinman, 2000; Craig, 2002; Hua et al., 2005), this study aimed to investigate the possibility of quantifying thermal interoception by means of a novel method based on the dynamic perception of temperature, named the *thermal matching task*. We believe that temperature perception could represent an ideal model to investigate interoception because it offers numerous advantages from the experimental and methodological point of view compared to other interoceptive modalities previously adopted in the field. First, stimulation can be experimentally controlled in the sense that we can systematically manipulate the temperature we deliver to the skin with a precision of ± 0.1-0.2°C (Somedic SenseLab AB, Hörby, Sweden). In contrast to pain or affective touch in which the affective facet can be prominent, temperature does not necessarily have a strong affective component when manipulated within the innocuous range (cool to warm perception), which is an advantage in experimental studies as it is easier to match conditions and raises fewer ethical issues than when administering pain. However, when the boundaries of the innocuous range are extended nearly to the pain threshold (cold to hot perception elicited by a temperature below 15°C or above 45°C, Kandel et al., 2000), these deviations are often accompanied by a strong affective state that signals individuals’ homeostatic needs and motivates us to take actions to re-establish homeostatic balance and optimal body functioning (e.g., change the temperature of the room or our clothing accordingly). This idea is congruent with the notion of thermosensation as an interoceptive modality that contributes to the perception of the physiological status of the body (Craig et al., 2000; Craig, 2002).

Thermoreception is a non-invasive way to investigate interoception compared to other modalities, such as gastric and bladder functions and pain. Traditionally, it has been argued that one of the advantages of focusing on the signals from the heart is that it is a constant signal in our life (Azzalini, Rebollo & Tallon-Baundry, 2019); our body and brain constantly receive cardiac signals, regardless of the extent to which we are aware of these signals. Thermosensation is similar in this respect, as our brain receives continuous signals about the temperature of the external environment from the receptors in the skin and about the internal temperature from receptors in the core of the body to maintain thermoneutrality, with the final aim of guaranteeing survival (Farrell et al., 2011, Tabarean, 2018). Consequently, as in the case of a rising heartbeat, we are prompted to pay attention to what is happening inside or outside our body as soon as there is a remarkable deviation from thermoneutrality. In other words, the body works as a thermostat, facilitating a constant dialogue between the inside and the outside of our body to maintain homeostasis. Thus, the body and the brain work in concert to maintain thermoneutrality, which is a task that involves our whole body (Proffitt, 2006; Davies et al., 2012).

Accordingly, the present study aimed to investigate thermosensation by means of two separate tasks: the well-established static temperature detection task, which targets cold and warm receptors, and a novel dynamic thermal matching task, which should elicit a combination of cold and warm receptors as well as CT receptors, given the use of dynamic stimulation as well as optimal thermal activation (Ackerley et al., 2014). We targeted temperatures within the thermoneutrality range (30-34°C, see Table 1) since we were more interested in exploring the interoceptive perception of temperature rather than the affective responses to such variations. We were particularly interested in individual differences in thermosensation, which have been recently reported to be specific for body parts (Luo et al., 2020). In line with previous studies in the context of affective touch and the CT system, we considered thermoreception in both hairy and non-hairy skin (see also Filingeri, Zhang, and Arens, 2018 for differences in thermosensation in hairy and non-hairy skin), since CT afferents have been found in significantly larger amounts in the hairy skin of the body (Nordin, 1990; Vallbo et al., 1993, 1999) and only sparsely in glabrous skin (Watkins, Dione, Ackerley, Backlund Wasling, Wessberg, & Löken, 2020). Recent evidence has shown optimal activation of CT afferents in response to slow, dynamic stimuli delivered at low pressure and neutral temperature (i.e., 32°C), as opposed to warmer or colder temperatures (Ackerley et al., 2014). We additionally targeted the involvement of the CT system in temperature perception by comparing participants’ performance on the dynamic thermal matching task, which supposedly activates the CT system (McGlone, Wessberg & Olausson, 2014), and on the static temperature detection task, which should not activate this system (but see Ackerley et al., 2018). Given the differences in thermal sensitivity between hairy and non-hairy skin (Filngeri et al., 2018), we also aimed to observe any potential differences in participants’ performance on the static and dynamic tasks, between the perception of cooling and warming, and between these two body locations.

A second aim of the present study was to explore the relationship between performance on thermal tasks and other interoceptive modalities, namely, heartbeat perception, affective touch, and pain detection (Heldestad et al., 2010; Craig, 2014 for a review). By comparing performance on these tasks targeting both internal and skin-mediated interoceptive signals, we aimed to address the open question of whether we can generalise interoceptive abilities or whether interoception comprises separated sub-modules, in which people show different and supposedly independent abilities. This is important because several accounts of interoception point towards the centrality of the posterior (e.g., Craig, 2003; Critchley et al., 2004) and anterior (e.g., Pollatos et al., 2007; Zaki, Davis, and Ochsner, 2012) insular cortex in processing interoceptive signals; however, it remains unclear whether the processing of interoceptive signals in the posterior insula and the subsequent integration of such signals in the anterior insula (Craig, 2009; Kuehn et al., 2016) are mirrored by a coherence in terms of interoceptive signal perception at the behavioural level (Ferentzi et al., 2018). In addition, cardiac interoception, as assessed by means of heartbeat detection tasks, is often used as a proxy for interoception more generally. However, this can be misleading, and we believe that a better understanding and a better description of the different interoceptive sub-modalities are both crucial for advancing the field.

In line with recent developments suggesting that interoception is a multidimensional construct (Garfinkel et al., 2015), for each interoceptive sub-modality, we distinguished between *interoceptive accuracy*, that is, the objective performance on an interoceptive task, i.e., perceptual detection or discrimination; *interoceptive sensibility*, which refers to subjective beliefs about the perception of bodily signals and is measured by means of self-reported questionnaires; and *interoceptive awareness*, or metacognitive awareness of interoceptive accuracy, which is one’s subjective confidence about the objective interoceptive performance that can be measured with confidence ratings (Garfinkel et al., 2015). This multi-dimensional manner of describing interoception allows us to capture the different facets of interoception - both at perceptual and cognitive levels - and to better describe the variability between individuals, which can enhance the understanding of interoception as a construct and highlight potential clinical implications of the present study (see Kalsha et al., 2018 for clinical implications of interoceptive research).

## Methods

### Participants

A total of sixty-four healthy participants (31 males and 33 females) were recruited using social media and advertising on the Karolinska Institutet campus. Two participants (one male and one female) were excluded because they did not meet the inclusion criteria; thus, a total of 62 participants were considered for data analysis. A priori power analysis based on previous studies in the field of interoception (Garfinkel et al., 2016; Crucianelli et al., 2018) suggested that a minimum sample of N = 62 provided enough power (92%) to detect our effects of interests (α = 0.05, effect size d = 0.4, two-tailed). Inclusion criteria were being 18-39 years old (mean age = 26.5 years, standard deviation = 5.4 years) and being right-handed. Exclusion criteria were having a history of any psychiatric or neurological conditions, taking any medications, having sensory or health conditions that might result in skin conditions (e.g., psoriasis), and having any scars or tattoos on the left forearm or hand. All participants were requested to wear short sleeves to make stimulation of the forearm easier. The study was approved by the Swedish Ethical Review Authority. All participants provided signed written consent, and they received a cinema ticket as compensation for their time. The study was conducted in accordance with the provisions of the Declaration of Helsinki 1975, as revised in 2008.

### Self-reported measures and interoceptive sensibility

Participants were asked to provide demographic information, such as age, weight and height (to calculate the body mass index, BMI), handedness and, for female subjects, the phase of the menstrual cycle at the time of testing (i.e., this is known to influence body temperature and consequently affect thermoregulatory processes; Kurz, 2008 for a review). Next, participants were asked to complete the following self-report questionnaires: the *Body Awareness Questionnaire* (BAQ), an 18-item questionnaire assessing body awareness (Shields, Mallory & Simon, 1989), and the *Body Perception Questionnaire* (very short form, BPQ), a 12-item questionnaire regarding perception of one’s body (Porges, 1993; Cabrera et al., 2017). The BAQ and BPQ were included as measures of interoceptive sensibility, that is, how aware participants reported being of their bodily sensations; the former questionnaire addresses more general body awareness, whereas the latter questionnaire targets bodily sensations more specifically, such as stomach and gut activity. Participants also completed the *Eating Disorder Examination Questionnaire* (EDE-Q 6.0, Fairburn & Beglin, 1994, 2008; Peterson et al., 2007) and the *Depression, Anxiety and Stress Scale – 21 Item* (DASS, Lovibond and Lovibond, 1995; Henry & Crawford, 2005). However, the BMI, EDE-Q and DASS were not considered in any of the following analyses because their inclusion lies beyond the scope of this manuscript.

### Interoceptive accuracy tasks

#### Heartbeat counting task (HCT)

The experimenter recorded the heartbeat frequency by means of a Biopac MP150 Heart Rate oximeter attached to the participant’s non-dominant index finger and connected to a Windows laptop with AcqKnowledge software (version 5.0), which enabled extraction of the actual number of heartbeats using the ‘count peak’ function. Care was taken to place the soft oximeter around the finger firmly but without being too tight to reduce the possibility that participants could perceive their pulse in their finger (Crucianelli et al., 2018; Murphy et al., 2019). As part of the task, a 5-minute heartbeat baseline was recorded to check for the presence of autonomic neuropathy. During this time, we presented the instructions for the heartbeat counting task (Schandry, 1981). Participants were asked to silently count their heartbeats between two verbal signals of ‘go’ and ‘stop’, without manually taking their pulse. Both of the participants’ hands were placed on the table to ensure that no body part was touched. Participants completed a practice trial of 15 seconds before proceeding to the three experimental trials (interval lengths of 25 s, 45 s, and 65 s), which were presented in a randomised order. Short breaks of 30 seconds were taken between each trial.

#### Temperature perception

##### (Dynamic) thermal matching task

Before proceeding with the task, the skin temperature of each participant’s palm and forearm was measured with a contactless thermometer (Microlife NC150) at three different locations at each site. These values are reported in Table 1 of the Supplementary Materials. This was done to control for any significant individual differences in skin temperature that could influence task performance. Then, participants were stroked with a 25×50 mm thermode attached to a thermal stimulator (Somedic MSA, SenseLab, Sweden) at reference temperatures of 30°, 32° or 34°C; these temperatures were within the range of neutral/innocuous temperatures so to mirror the performance at the heartbeat counting task, which is usually performed at rest. Participants were instructed to pay close attention to this reference temperature because their task would be to match it by verbally indicating whenever they felt the same temperature again. Next, in each experimental trial, the experimenter touched the participant with the thermode set at different temperatures starting from ± 8°C (which is 25% of the neutral temperature of 32°C; whether the starting temperature was +8°C or −8°C from the reference temperature was counterbalanced across participants) of the reference temperature (range 22-38°C for the reference temperature of 30°C; range 24-40°C for the reference temperature of 32°C; range 26-42°C for the reference temperature of 34°C). The task followed a staircase procedure, that is, the temperature was either increased (i.e., from cool to warm) or decreased (i.e., from warm to cool) towards the reference temperature in discrete steps of 2°C. Temperature was increased or decreased until participants verbally indicated they felt the reference temperature or until the maximum or minimum temperature was reached (± 8°C from the reference temperature, opposing the starting temperature) for a total of 9 potential strokes per trial, with a break of 3 seconds between trials. Participants were instructed to try to match the reference temperature that they previously experienced. The correct answer was always the reference temperature, and the order in which the reference temperatures were presented as well as the order of increasing and decreasing trials varied across trials to avoid anchor effects of the initial values (e.g., if one participant started with increasing trials based on one reference temperature, then they would start with decreasing trials for the following reference temperature, see Tajadura-Jiménez et al., 2015 for a similar approach in an embodiment paradigm). Two trials per reference temperature were repeated, one increasing and one decreasing, for a total of 6 trials presented in randomised order. The duration of each stroke was kept constant at 3 seconds; the velocity of tactile stimulation was CT-optimal (3 cm/s) and the direction of movement was always proximal to distal with respect to the participant. No additional pressure was applied aside from the weight of the thermode. The same procedure was repeated on the outer forearm (hairy skin) and on the palm (non-hairy skin) in areas of 9×4 cm.

##### (Static) temperature detection task

As in the dynamic thermal matching task, the tactile stimulus was delivered using the Somedic MSA Thermal Stimulator. The detection of cold and warm static thermal stimuli was measured by means of the well-established Martsock methods of the limits (Fruhstorfer et al., 1976), and we used the same protocol adopted by Heldestad et al., 2010. The experimenter held the thermode on the area of interest (left forearm or palm) without applying any additional pressure. The thermode was not secured on the forearm or hand to avoid any additional tactile signals that could interfere with the detection of temperature. Participants were asked to hold a response button using their right hand and to press it as soon as they perceived a change in temperature of any kind (i.e., warmer or colder than the previous perceived temperature, Heldestad et al., 2010). The starting temperature was always neutral (32°C); the maximum probe temperature was set to 50°C, and the minimum was set to 10°C for safety reasons. As soon as the button was pressed, the temperature automatically changed in the opposite direction and returned to the baseline temperature of 32°C; the temperature stayed at 32°C for 5 seconds before moving to the next trial. The temperature changed at a rate of 1°C/s and returned to baseline at a speed of 4°C/s. This method has been widely used to detect neuropathy in clinical settings, and it includes a total of five warm and five cold trials, presented in two blocks (warm and cold blocks). The procedure was repeated twice: once on the left forearm and once on the left palm, in a randomised order.

#### Affective touch task

This task takes advantage of the discovery that affective, hedonic touch on the skin can be reliably elicited by soft, light stroking at specific velocities within the range of 1-10 cm/s that activate a specialised peripheral system of C-tactile afferents (Löken, Wessberg, Morrison, McGlone & Olausson, 2009; McGlone, Valbo, Olausson, Löken, & Wessberg, 2007). Touches were delivered using a soft brush (i.e., precision cheek brush No 032, Åhléns, Sweden) on the left forearm (hairy skin that contains CT afferents) and left palm (non-hairy skin, where CT afferent activity has only partially been reported), and the task of the participants was always to verbally rate the pleasantness of the touch using the rating scale. Touches were delivered at seven velocities (0.3, 1, 3, 6, 9, 18 and 27 cm/s). Two slow velocities of 3 and 6 cm/s are typically perceived as more pleasant (i.e., CT optimal velocities) compared to the borderline optimal velocities (1 and 9 cm/s) and the CT non-optimal speeds (0.3, 18 and 27 cm/s, Löken et al, 2009). Each velocity was presented three times, for a total of 21 stroking trials per location (palm and forearm, in randomised order) and the direction of movement was always proximal to distal with respect to the participant.

#### (Static) pain detection task

The procedure of this task followed the same protocol to detect thermal pain thresholds used by Heldestad et al., 2010, and it was similar to the one described for static temperature detection. However, here, participants were instructed to press the button as soon as they perceived that the thermal stimulation was becoming uncomfortable or painful (Heldestad et al., 2010). When providing the instructions, the experimenter clarified that the task was to press the button as soon as the sensation of discomfort or pain was beginning (i.e., detection) rather than when the pain was unbearable (i.e., threshold). We performed the procedure in the left palm (non-hairy) and forearm (hairy), and we tested pain detection following warm stimuli only for a total of five trials per location. The baseline starting temperature was 32°C, and the maximum temperature was 50°C for safety reasons. The temperature changed at a rate of 2°C/s, whereas the return to baseline in all tests occurred at a speed of 4°C/s.

### Interoceptive metacognitive awareness: Confidence and prior beliefs

In line with recent models of interoception (Garfinkel et al., 2015; 2016), we also measured metacognitive awareness in relation to interoception. We collected information about this measure as confidence after each answer (i.e., online) and as prior belief before participants completed each task (i.e., offline); these data have been analysed separately (Fleming et al., 2016). After receiving the instructions about each task and having been given the opportunity to ask any questions they might have, participants were asked to provide a prospective estimation of their ability to successfully complete the task by means of a rating scale ranging from 0 (not at all accurate/total guess) to 100 (very accurate) (Beck et al., 2019). Furthermore, participants were also asked after each individual trial within the tasks to rate their confidence with their answers (as in Garfinkel et al., 2015; Beck et al., 2019). This confidence rating was chosen on an 11-point scale ranging from 0 (not at all) to 10 (extremely). This was done for each trial of the tasks except for the static temperature detection task and static pain detection task, as these followed a standardised method-of-limits procedure, whereby temperature changes in a continuous manner; providing individual confidence ratings after each trial during the task would have disrupted the actual performance.

### Experimental procedure

Participants were welcomed into the experimental room, and they were asked to sit on a table opposite the experimenter. Upon arrival, they were asked to sign a consent form and to complete the following questionnaires presented in an online format: the demographic questionnaire, BAQ, BPQ, EDE-Q and DASS. The questionnaires were always presented at the beginning of the experimental procedure to ensure that participants were given some time to stay at rest before completing the *heartbeat counting task*, which was the first interoceptive task that all participants completed. Given that previous studies showed that the heartbeat counting task might be influenced by other activities (e.g., Ring et al., 2015; Brener and Ring, 2016), we decided to conduct this task first (for an overview of procedures and tasks, see Figure 1 and Table 2). Participants were given the choice to either keep their eyes closed or open, whichever helped them feel more comfortable, in order to be as accurate as possible. The aforementioned experimental procedure prior to the thermal matching task took approximately 30 minutes, giving participants the opportunity to acclimatise themselves before proceeding with the dynamic *thermal matching task*. Participants were asked to wear a disposable blindfold and to place their left arm on the table to complete the dynamic *thermal matching task*, following the method fully described in Method section above. Participants were asked to pay close attention to each reference temperature because they were given the possibility to feel it just once. Upon completion, participants removed the blindfold, and they were given a short break before beginning the *affective touch task*. As part of this task, they were familiarised with the pleasantness rating scale, and the experimenter identified and marked two identical areas of 9×4 cm on the left forearm and palm with a washable marker, as was done in previous studies (Crucianelli et al., 2013; 2016; 2018). This was performed to control the stimulated area and the pressure applied during the touch by checking that the tactile stimulation was applied just inside the marked areas (more pressure would result in a wider spreading of the brush, that is, the tactile stimulation would be applied outside of the marked borders). Alternating the stimulated areas would counteract the fatigue of the CT fibres (McGlone et al. 2012). Participants were asked to wear the blindfold again for the entire duration of the affective touch task. Next, participants could take a break from wearing the blindfold before starting the *static temperature detection task*. No break was taken between the cold and warm blocks, but participants were only allowed to remove the blindfold at the very end of the task. The last part of the experimental procedure consisted of the *static pain detection task*, for which participants were asked to wear the blindfold once again. All the experimental tasks were conducted on the left, non-dominant hand or forearm. The starting location for each task was alternated between the forearm and the palm (e.g., participants starting one task on the palm next completed the task with the forearm; those who started one task with the forearm completed the following task with the palm). The order of the tasks was kept constant (with internal randomisation) (Figure 1 and Table 2). The pain detection task was performed last as to not arouse the body or cause hypoalgesia, which could affect performance on the other tasks (Gröne et al., 2012). The entire experimental procedure lasted approximately one hour, and participants were offered a wipe to remove the marker from the skin and were provided with a full debriefing at the end of the session. Testing took place in a testing room with constant temperature and humidity, with no significant changes in temperature between the beginning (M = 22.55°C, SD = 0.49) and the end (M = 23.10°C, SD = 0.47) of the testing session.

**Table 2.** Overview of the structure of the different tasks. The basic tasks are described in the order that they were completed during the experimental procedure. Each row represents a task, and each column describes one step of each task, starting with pre-task estimation (or prior belief of performance), followed by the name of the actual task with its description in terms of method (number of trials and body sites) and outcome measures (interoceptive accuracy). The final column refers to the participants’ performance confidence, the data of which were collected after each trial of the heartbeat counting task, dynamic thermal matching task and affective touch task only.

| Interoceptive modality | Pre-task estimation | Interoceptive task | Task description | Outcome measures (interoceptive accuracy) | Post-trial confidence |
| --- | --- | --- | --- | --- | --- |
| Questionnaires | N/A | Interoceptive sensibility | Body Awareness Questionnaire<br>Body Perception Questionnaire | Values between 18 - 126<br>Values between 12 - 60 | N/A |
| Cardiac | How well are you going to perform on this task? (0, not well at all - 100, very well) | Heartbeat counting task | 3 trials (25 s, 45 s, and 65 s) | Values between 0 - 1 | How confident are you with your answer? (0, not at all - 10, very) |
| Dynamic temperature |  | Thermal matching task | 3 temperatures (30, 32, and 34°C)<br>2 body locations (palm and forearm)<br>2 staircase approaches per temperature (increasing and decreasing temperature) | Values between 0 - 1 |  |
| Tactile affectivity |  | Affective touch task | 7 velocities (0.3, 1, 3, 6, 9, 18, 27 cm/s)<br>2 body locations (palm and forearm) | Pleasantness of touch 0-100 & tactile sensitivity (CT – non-CT) |  |
| Static temperature |  | Temperature detection | Method of limit<br>5 trials for warm and 5 trials for cold | Detection temperature and standard deviations | N/A |
| Static pain |  | Pain detection | Method of limit<br>5 trails for warm pain | Detection temperature and standard deviations |  |

**Figure 1.**
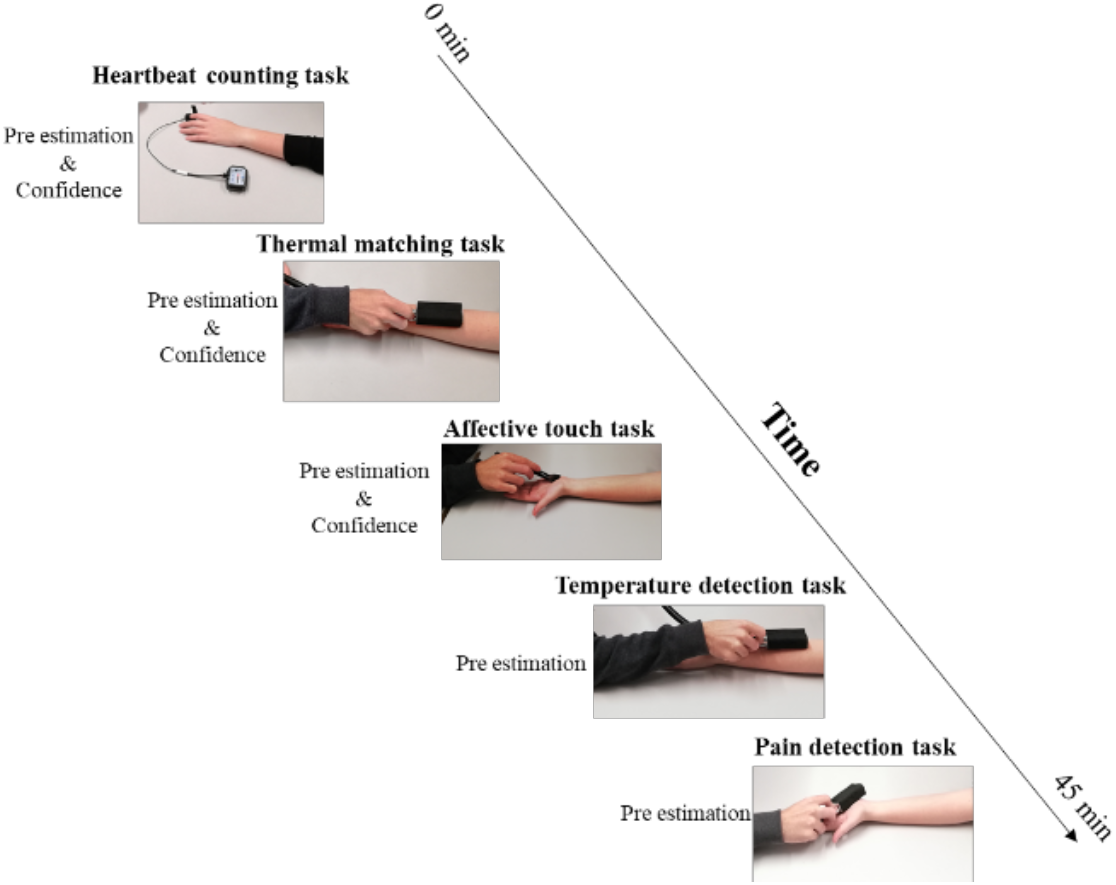
The experimental procedure. The heartbeat counting task was conducted using the BioNomadix system of a wearable wireless device connected to a Biopac MP150 system. The thermal matching task, temperature detection task and pain detection task were conducted using the thermode connected to the Somedic thermal stimulator. In the affective touch task, tactile stimulation was delivered with a soft brush. All the tasks were repeated on the forearm and on the palm in a randomised order.

### Design and plan of analysis

All data were analysed with the Statistical Package for Social Sciences (SPSS), version 26. The data were tested for normality by means of the Shapiro-Wilk test and were found to be non-normal (p < .05). Subsequent two-step approach transformations (Templeton, 2011) did correct for the normality violations; therefore, parametric tests were used to analyse the data (described below). The false discovery rate (FDR, Benjamini & Hochberg, 1995) was used to correct for multiple correlations (we reported the corrected values for the significant effects); this method is widely used when a large number of multiple comparisons is applied and it controls the proportions of false rejections out of all rejections (Benjamini, 2010). Bonferroni-corrected post hoc comparisons were used to follow up significant effects and interactions. All *p* values are 2-tailed unless otherwise specified.

First, we focused on the analysis of each task separately. As in Garfinkel et al., 2016, we first assessed whether there was a relationship between the dimensions of interoception (accuracy, confidence and prior beliefs) for each modality separately (cardiac, dynamic and static temperature, affective touch, and pain). Then, we investigated the relationship between the different interoceptive modalities and dimensions. Specifically, we ran correlational analyses to investigate the relationship between accuracy and confidence across the modalities. In secondary analyses using parametric correlational analyses, we also explored the relationship between accuracy and prior beliefs of performance, as well as the relationships between the interoceptive dimensions and individual differences in the questionnaires probing self-reported interoceptive awareness and bodily awareness (interoceptive sensibility). The results of secondary analyses are reported in Supplementary Materials only, for brevity.

We also performed Bayesian correlations for our main analyses of interest (i.e., correlations between accuracy - objective performance - across different interoceptive modalities). Bayesian correlations produce a Bayes factor (BF) as the main output index. BF_01_ indicates the probability supporting the null over the alternative hypotheses (e.g., a BF_01_ = 8 means that H_0_ is 8 times more likely to be true than H_1_). By convention, BFs between 0.33 and 3 are considered inconclusive (Jarosz & Wiley, 2014; Lee and Wagenmakers, 2014; Biel and Friedrich, 2018).

#### Interoceptive accuracy

We calculated the cardiac interoceptive accuracy (*heartbeat counting task)* by means of the following standardised formula that allowed us to compare the counted and recorded heartbeats (Shandry, 1981):

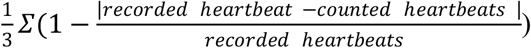

For the other tasks, the focus was 1) to explore whether there was a significant effect of touch location (hairy vs. non-hairy skin) and 2) to obtain an accuracy value that could resemble, and therefore be compared to, the interoceptive accuracy measured by means of the heartbeat counting task. This was done to ensure that levels of accuracy were equated across the modalities.

For the dynamic *thermal matching task*, we used the following formula:

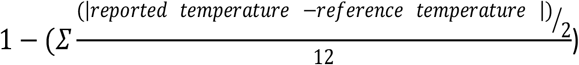

where 12 represents the total number of options presented to participants (regardless of direction - overestimation or underestimation of temperature) across the three trials. Both of these formulas provide a value between 0 and 1, with 0 suggesting poor performance and 1 indicating optimal performance on the task. We kept the order of the increasing and decreasing stimuli separate given the different mechanisms and skin responses known to be involved when perceiving cooling and warming stimuli (Nordin, 1990; Wiklund Fernström & Wessberg, 2003; Olausson, Wessberg, Morrison, McGlone, & Vallbo, 2010, for a review). Thus, for each subject, we obtained one increasing and one decreasing accuracy value for the forearm and for the palm. We provide an additional control analysis that focused on the perception of the three temperatures separately (30, 32 and 34°C) in hairy and non-hairy skin in the Supplementary Materials.

The *affective touch task* was analysed as in previous studies (e.g., Crucianelli et al., 2018). We obtained the scores for pleasantness for the CT-optimal, borderline and CT-non-optimal velocities by averaging the scores of tactile pleasantness in each of these categories. This allowed us to investigate the main effect of velocity and skin site on pleasantness by means of a repeated measures ANOVA. For the purpose of this study, our main variable of interest was the so-called ‘affective touch sensitivity’ (Crucianelli et al., 2018; Kirsch et al., 2020), which describes the individual’s ability to differentiate levels of pleasantness between affective and neutral touch, without taking into account the total pleasantness. Thus, we averaged the pleasantness scores for CT-optimal velocities and for CT-non-optimal velocities, and we calculated the differences between these two measurements to obtain one tactile sensitivity score for the forearm and one for the palm. This differential score was then used in the analysis to investigate the relationship with participants’ performance on the other interoceptive tasks.

Next, for both *static temperature detection* and *static pain detection*, we were interested in both the sensitivity (i.e., the smallest change in temperature a person could detect) and the consistency or precision (i.e., the variability in the individual responses across the different trials, quantified as standard deviations) of the detection across trials. As a proxy of interoceptive accuracy, we calculated the relationship between sensitivity and consistency and obtained one detection accuracy value for cold temperature, one value for warm temperature and one value for warm pain for both hairy and non-hairy skin using the following formula:

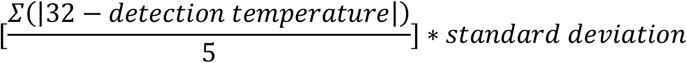

where 32°C is the baseline starting temperature, detection temperature is the temperature that participants recognise as different (warmer or cooler) from baseline, and 5 is the total number of trials; we multiplied by the standard deviation to account for the individual variability in responses across trials. We developed this formula to take into account both the accuracy (i.e., how many degrees are necessary for the participants to detect a change) and the precision (i.e., how consistent participants are in their performance across trials). We then used these detection accuracy values to investigate the relationship with the other interoceptive modalities.

#### Interoceptive metacognitive awareness: Confidence and prior beliefs

In terms of metacognitive interoception, we focused both on ‘offline’ insight into participants’ own abilities *before* they completed the tasks and on ‘online’ confidence in their own answers reported immediately *after* each trial of the heartbeat counting task, dynamic thermal matching task and affective touch task. Specifically, metacognitive awareness for each interoceptive modality was operationalised as the extent to which pre-estimation of performance on each task and confidence predicted accuracy (Garfinkel et al., 2015; 2016). This was analysed by means of multiple regressions, with pre-estimation and confidence as the main predictors and accuracy as the outcome variable. The offline metacognitive measure was computed separately for cardiac, dynamic thermal matching task, affective touch, static temperature and static pain detection responses to provide five measures of metacognitive awareness. The online metacognitive measure (i.e., confidence) was obtained only for the cardiac interoception, dynamic thermal matching, and affective touch tasks because the static temperature detection task and static pain detection task followed a standardised method-of-limits procedure; providing individual confidence ratings after each trial during the task would have disrupted the participants’ actual performance. Confidence ratings were averaged over trials for all the tasks.

## Results

### Demographics and interoceptive sensibility

The mean scores and standard deviations for BMI, interoceptive sensibility (as measured by means of the BAQ and BPQ), EDE-Q and DASS scores are reported in Table 3. No effect of sex on any of these measures was found, except for the EDE-Q.

**Table 3.** Mean and standard deviations for the participants’ body mass index (BMI); Body Awareness Questionnaire (BAQ) scores; Body Perception Questionnaire (BPQ) scores; Eating Disorders Examination Questionnaire (EDE-Q) scores; and the scores on the depression, anxiety and stress scales.

|  | BMI | BAQ | BPQ | EDE-Q | Depression | Anxiety | Stress |
| --- | --- | --- | --- | --- | --- | --- | --- |
| **Total sample** | 22.87 (2.99) | 79.79 (18.38) | 29.90 (8.77) | 1.42 (1.12) | 4.13 (3.87) | 4.02 (3.52) | 5.76 (4.05) |
| **Females (32)** | 22.19 (2.90) | 79.66 (19.86) | 30.00 (9.22) | 1.70 (1.14) | 4.16 (4.05) | 3.38 (2.79) | 5.88 (4.14) |
| **Males (30)** | 23.60 (2.95) | 79.93 (16.99) | 29.80 (8.41) | 1.11 (1.04) | 4.10 (3.74) | 4.70 (4.10) | 5.63 (4.02) |
| ***F* (*p*) values** | 3.62 (0.06) | 0.03 (0.95) | 0.08 (0.93) | 4.56 (0.04)* | 0.03 (0.96) | 2.23 (0.14) | 0.05 (0.82) |

### Interoceptive accuracy across modalities

#### Heartbeat detection task

The mean cardiac interoceptive accuracy score was 0.64 (SD = 0.25) in the present sample. This value is in line with those reported in previous studies (e.g., Tsakiris, Jiménez & Costantini, 2011; Crucianelli et al., 2018). The mean confidence score was 5.77 (SD = 2.28). One-way ANOVA revealed no effect of sex on cardiac accuracy (*F* (1, 61) = 0.128, *p* = 0.722, ηp^2^ = 0.002).

#### (Dynamic) thermal matching task

As mentioned in the Methods section, we obtained one increasing (staircase) temperature accuracy value for the forearm and one for the palm and one decreasing (staircase) value for the forearm and one value for the palm. The results of the 2 (location: palm vs. forearm) x 2 (staircase: increasing vs. decreasing) repeated measure ANOVA revealed a significant main effect of location (*F* (1, 61) = 5.00, *p* = 0.029, η_p_^2^ = 0.084, Figure 2), with participants being more accurate in the detection of dynamic temperature in the forearm (M = 0.81; SD = 0.16) than in the palm (M = 0.76; SD = 0.17). No main effect of staircase (*F* (1, 61) = 0.142; *p* = 0.707, η_p_^2^ = 0.002) or significant interaction (*F* (1, 61) = 0.299; *p* = 0.586, η_p_^2^ = 0.005) was found. No effect of sex was found for any of the variables of interest as investigated by means of separate one-way ANOVAs (all *F*_s_ between 0.024 and 0.467; all *p*_s_ between 0.497 and 0.878); thus, sex was not considered in subsequent analyses.

**Figure 2.**
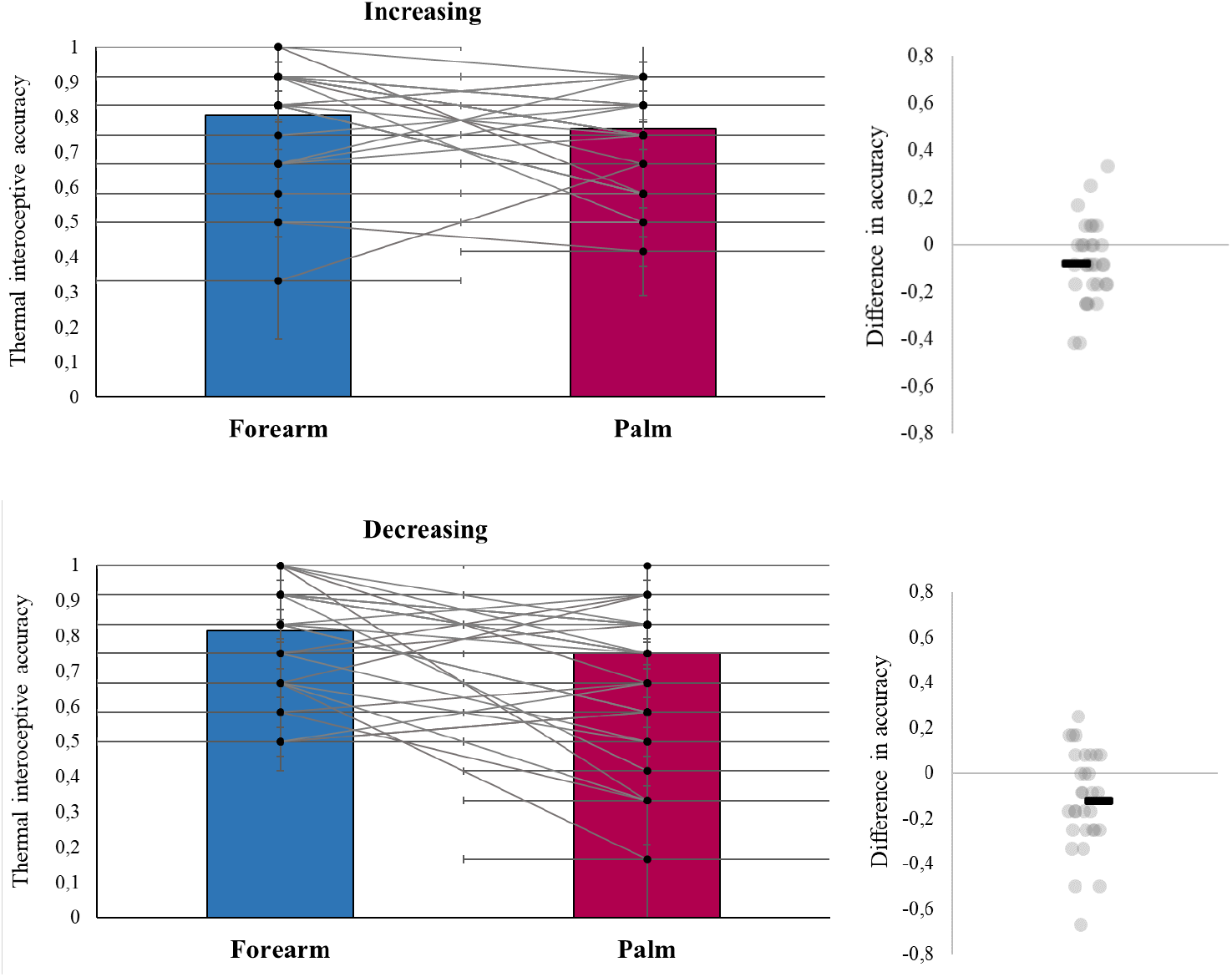
Mean, data distribution and standard errors for the dynamic thermal matching task, performed on the forearms (in blue) and palms (in burgundy) of the participants. The panel on top shows task performance during increasing trials, and the panel on the bottom shows task performance on decreasing trials. The graphs on the right-hand side represent the distribution of the differences between task performance regarding the forearm and palm for each participant.

#### Affective touch task

We averaged the CT optimal velocities (3 and 6 cm/s), borderline velocities (1 and 9 cm/s) and the CT-non-optimal velocities (0.3, 18 and 27 cm/s) to obtain three velocity variables. As expected, there was a main effect of velocity on touch pleasantness (*F* (2, 122) = 40.07, *p* < 0.001, ηp^2^ = 0.417). Bonferroni corrected post hoc analysis (α = 0.017) revealed that slow, CT-optimal touch was rated as more pleasant (M = 58.24; SD = 23.02) than fast, CT-non-optimal touch (M = 46.08; SD = 22.23; *t*(61) = 6.67; *p* < 0.001, Figure 3, Figure 1 in Supplementary Materials) and touch delivered at borderline velocities (M = 55.91; SD = 22.87; *t*(61) = 2.77; *p* < 0.001). There was also a significant difference between borderline and CT-non-optimal touch (*t*(61) = 6.47; *p* < 0.001). There was a main effect of location (*F* (1, 61) = 5.708, *p* = 0.020, ηp2 = 0.092), with touch being rated overall as more pleasant in the forearm (M = 55.32; SD = 22.61) than in the palm (M = 51.50; SD = 23.80), consistent with previous findings (Löken, Evert and Wessberg, 2011). There was a significant interaction between velocity and location (*F* (2, 122) = 4.896; *p* = 0.009, ηp2 = 0.080). Bonferroni corrected post hoc analysis (α = 0.017) revealed a significant difference between the forearm and palm only in the perception of slow, CT-optimal touch *t*(61) = −2.93; *p* = 0.005), but not in the perception of borderline (*t*(61) = −1.19; *p* = 0.241) or CT-non-optimal touch (*t*(61) = −0.46; *p* = 0.650). No effect of sex on any of the touch pleasantness scores was found (all *F*_s_ between 0.002 and 2.394; all *p*_s_ between 0.127 and 0.966).

**Figure 3.**
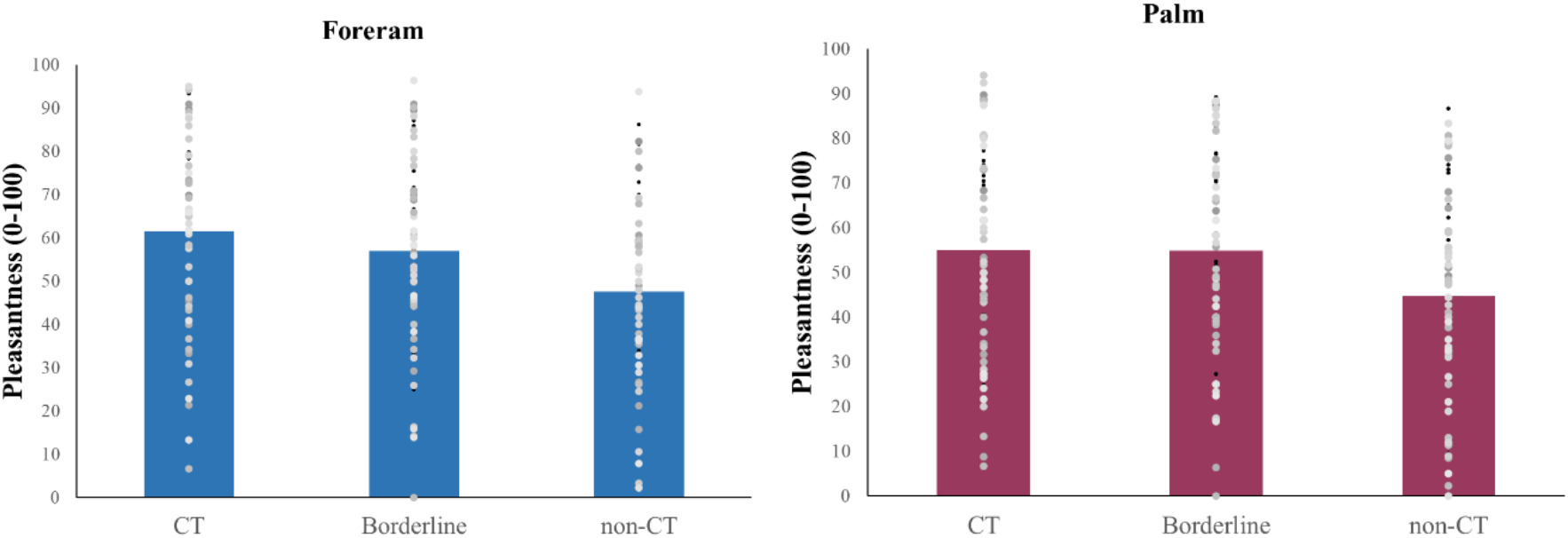
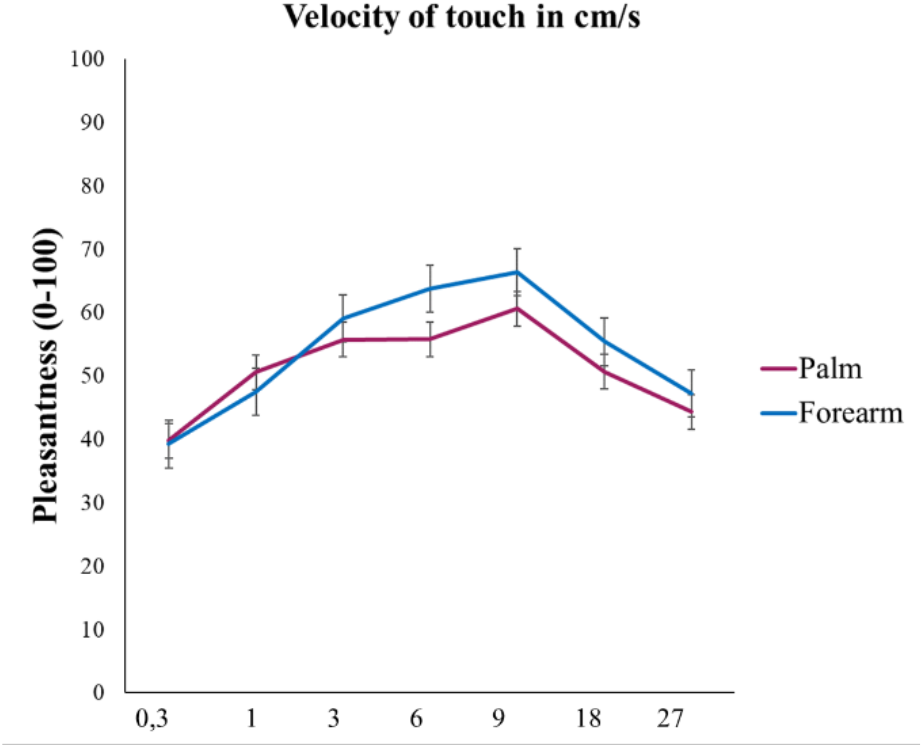
**Top panels:** Mean and data distribution for the pleasantness scores for CT-optimal, borderline and CT-non-optimal velocities for the forearm and palm. **Left panel:** Mean and standard errors for the same dataset of the affective touch task for each velocity (velocities are reported in cm/s), showing the main effect of velocity and location (forearm vs. palm) on touch pleasantness. A full account of the mean pleasantness for each velocity is reported in Figure 1 of the supplementary materials.

#### (Static) temperature detection task

We compared the smallest change in temperature participants could detect when temperature was increasing (warm) or decreasing (cool) from the neutral starting temperature of 32°C in both hairy (forearm) and non-hairy skin (palm). The results of the 2 (temperature: warm vs. cool) × 2 (location: forearm vs. palm) repeated measures ANOVA revealed a main effect of temperature (*F* (1, 61) = 67.74; *p* < 0.001, ηp^2^ = 0.855), suggesting that participants could detect cooling (M = 1.75°C; SD = 1.08) quicker or with a significantly smaller change in temperature compared to warming (M = 2.63°C; SD = 1.36, Figure 4). There was a non-significant main effect of location (*F* (1, 61) = 3.71; *p* = 0.06, ηp^2^ = 0.066), with participants needing a smaller but non-significant variation in terms of °C to detect changes in temperature in the forearm (M = 2.05°C; SD = 1.25) compared to the palm (M = 2.34°C; SD = 1.35, Figure 4). There was a significant interaction between temperature and skin site (*F* (1, 61) = 5.90; *p* = 0.02, ηp^2^ = 0.060). Bonferroni-corrected post hoc analysis (α = 0.025) revealed a significant difference between hairy and non-hairy skin in the detection of cold temperatures (*t*(61) = −3.47, *p* < 0.01) but not warm temperatures (*t*(61) = −0.18, *p* = 0.86, see Figure 4).

**Figure 4.**
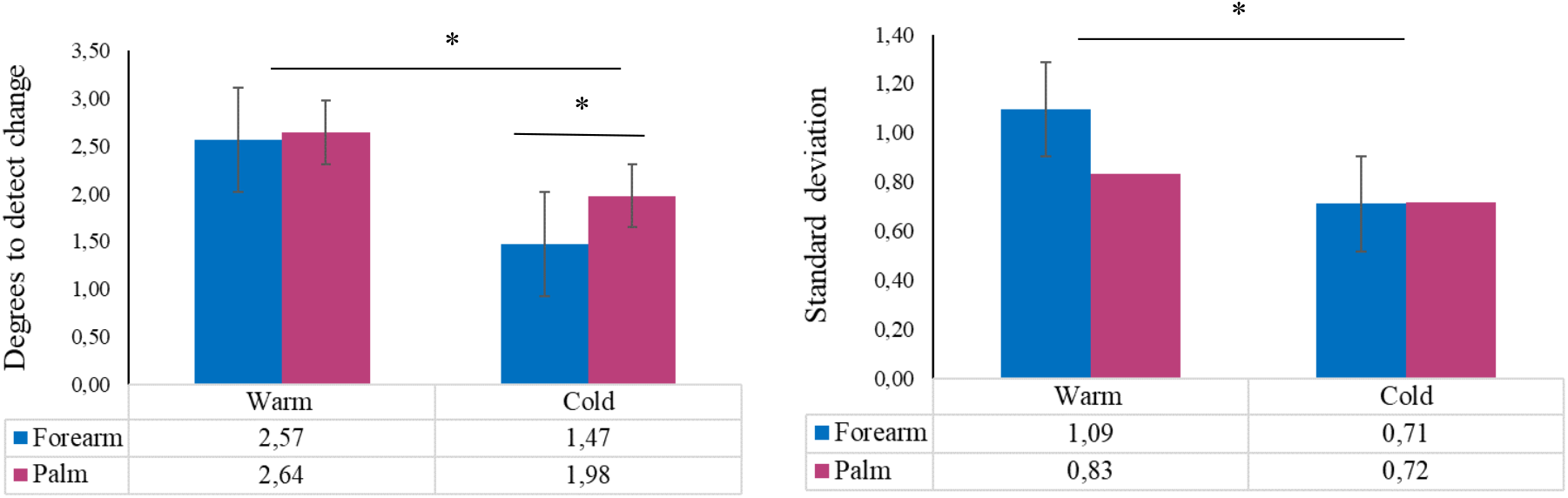
Means and standard errors for the static temperature detection task, showing the main effect of temperature on participants’ ability to detect temperature (quantified as degrees to detect the change, measured in °C) and consistency in detecting changes in temperature for warm and cool temperatures (quantified as standard deviation). * indicates significant differences, *p* < 0.05.

There was a trend towards significance for the effect of sex on participants’ temperature detection of both warm (*F* (1, 61) = 3.97; *p* = 0.051) and cold temperatures (*F* (1, 61) = 3.88; *p* = 0.053) on the palm, but not on the forearm (warm: *F* (1, 61) = 2.56; *p* = 0.115; cold: *F* (1, 61) = 0.05, *p* = 0.831). That is, female participants could detect changes in temperature more promptly than male participants on the palm (female: M = 2.03, SD = 0.85; male: M = 2.66, SD = 1.61) but not on the forearm (female: M = 1.94; SD = 0.96; male: M = 2.15; SD = 1.20).

In terms of consistency (operationalised as the standard deviation) in the perception of thermal static stimuli, the results of the 2 (temperature: cool vs. warm) × 2 (location: forearm vs. palm) repeated measures ANOVA revealed a main effect of temperature (*F* (1, 61) = 7.09; *p* = 0.01, ηp^2^ = 0.104), suggesting that participants were more consistent in the detection of cold temperatures (M = 0.71; SD = 0.65) than warm temperatures (M = 0.96; SD = 0.68, Figure 4). No significant main effect of location (*F* (1, 61) = 1.19; *p* = 0.28, ηp^2^ = 0.019) or interaction (*F* (1, 61) = 2.16; *p* = 0.15, ηp^2^ = 0.034) was found. There was an effect of sex on the consistency in the detection of warm temperatures in both the palm (*F* (1, 61) = 6.514; *p* = 0.013) and forearm (*F* (1, 61) = 5.041; *p* = 0.028) but not for the detection of cold temperatures (palm: *F* (1, 61) = 0.094; *p* = 0.760; forearm: *F* (1, 61) = 0.018; *p* = 0.893). That is, female participants were significantly more consistent than male participants in the detection of static warming in the palm (female: M = 0.59; SD = 0.37; male: M = 1.09; SD = 1.04) and forearm (female: M = 0.84; SD = 0.74; male: M = 1.37; SD = 1.08).

#### (Static) pain detection task

Two paired sample t-tests were used to investigate differences between hairy (forearm) and non-hairy (palm) skin in the temperature necessary for participants to detect pain and the consistency (i.e., standard deviations) in reporting pain sensation. The results showed no significant main effects of body site on individual thresholds (*t*(61) = −1.12; *p* = 0.27; forearm, M = 42.26; SD = 4.47; palm, M = 42.72; SD = 4.50) or on consistency in detection (*t*(56) = −0.70; *p* = 0.49, see Figure 2 in Supplementary Materials). No effect of sex on pain detection was found (all *F*_s_ between 0.139 and 3.06; all *p*_s_ between 0.086 and 0.711).

#### Relationship across interoceptive modalities

First, we focused on the relationship between performance on the heartbeat counting task and other interoceptive modalities, as in Garfinkel et al. 2016. Cardiac interoceptive accuracy was significantly related to the dynamic thermal matching task on the forearm in the decreasing (cooling) condition only (*r* = 0.27, *p* = 0.03, BF_01_ = 0.62; Figure 5) and not in the increasing (warming) condition (*r* = 0.06; *p* = 0.61; BF_01_ = 9.47). However, the relationship between decreasing thermal accuracy on the forearm and cardiac accuracy was not significant after applying FDR (Benjamini-Hochberg adjusted *p* value = 0.18).

No significant relationship was found between performance on the heartbeat counting task and the dynamic thermal matching task on the palm (increasing: *r* = 0.03, *p* = 0.79, BF_01_ = 9.67; decreasing: *r* = 0.16, *p* = 0.22, BF_01_ = 6.31).

**Figure 5.**
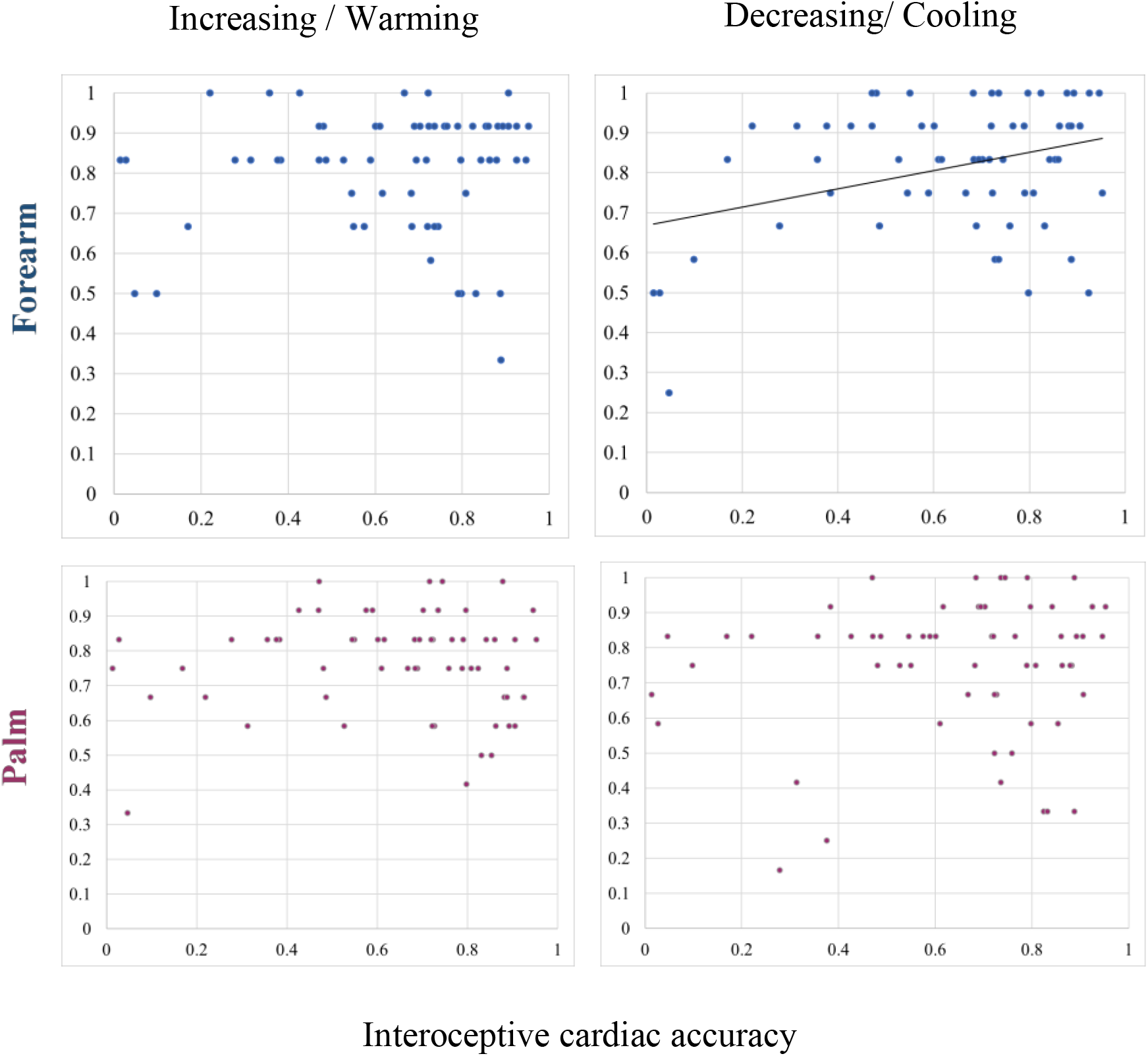
Scatter plot representing the relationship between performance on the heartbeat counting task (cardiac accuracy) on the x-axis and performance on the thermal matching task on the forearm only when the temperature was presented in decreasing staircases (i.e., cooling). The correlation was not significant after applying FDR corrections. Both the x- and y-axes refer to accuracy, which ranges from 0 to 1 and is derived using the formulas specified in section 2.4.

BF_01_ indicated that the null hypothesis (cardiac accuracy not related to thermal accuracy) was more likely than the alternative hypothesis (cardiac accuracy related to thermal accuracy) (BF_01_> 1), except for the relationship between cardiac accuracy and the thermal matching task in the forearm when temperature was decreasing (BF_01_< 1). However, a BF between 0.33 and 3 is usually considered inconclusive (Lee and Wagenmakers, 2014; Biel and Friedrich, 2018).

In line with previous findings (Crucianelli et al., 2018), performance on the heartbeat counting task was not related to affective touch sensitivity, that is, the difference in pleasantness between slow and fast touch on the forearm (*r* = −0.07, *p* = 0.59, BF_01_ = 7.38) or on the palm (*r* = −0.08, *p* = 0.54, BF_01_ = 7.64, see Figure 3 in Supplementary Materials). BF_01_ indicated that the null hypothesis (cardiac accuracy not related to tactile sensitivity) was 7.38 (for the forearm) and 7.64 (for the palm) times more likely than the alternative hypothesis (cardiac accuracy related to tactile sensitivity) (BF_01_> 1).

Cardiac interoceptive accuracy was not significantly related to the detection of static temperature on the forearm (see Table 4) or on the palm (see Table 5). BF_01_ indicated that the null hypothesis (cardiac accuracy not related to static temperature detection) was more likely than an alternative hypothesis (cardiac accuracy related to static temperature detection) (all BF_01_> 1). Finally, no significant relationship was found between cardiac interoceptive accuracy and (warm) pain detection (see Table 4 for forearm data and Table 5 for palm data). BF_01_ indicated that the null hypothesis (cardiac accuracy not related to pain detection) was more likely than an alternative hypothesis (cardiac accuracy related to pain detection) (all BF_01_> 1).

**Table 4.** Correlational matrix describing the relationship between performance on the different interoceptive tasks (i.e., interoceptive accuracy) on the forearm. Thermal interoceptive accuracies when the temperature is decreasing (cooling) and increasing (warming) are significantly correlated. *P* values correspond to original FDR corrected values.

| Forearm |  | Heartbeat counting task | Thermal matching task |  | Affective touch task | Temperature detection |  | Pain detection |
| --- | --- | --- | --- | --- | --- | --- | --- | --- |
|  |  |  | Increasing | Decreasing |  | Warm | Cold |  |
| Heartbeat counting task |  | 1 |  |  |  |  |  |  |
| Thermal matching task | Increasing | $r = 0.06$<br>$p = 0.980$<br>$BF_{01} = 9.47$ | 1 | | | | | |
| | Decreasing | $r = 0.27$<br>$p = 0.180$<br>$BF_{01} = 0.62$ | $r = 0.36$<br>$p = 0.180$<br>$BF_{01} = 0.02$ | 1 | | | | |
| Affective touch task | | $r = -0.09$<br>$p = 0.935$<br>$BF_{01} = 7.38$ | $r = 0.05$<br>$p = 0.852$<br>$BF_{01} = 9.99$ | $r = 0.06$<br>$p = 0.980$<br>$BF_{01} = 8.58$ | 1 | | | |
| Temperature detection | Warm | $r = -0.14$<br>$p = 0.840$<br>$BF_{01} = 5.66$ | $r = -0.13$<br>$p = 0.840$<br>$BF_{01} = 5.07$ | $r = -0.05$<br>$p = 0.852$<br>$BF_{01} = 9.99$ | $r = -0.06$<br>$p = 0.892$<br>$BF_{01} = 6.19$ | 1 | | |
| | Cold | $r = 0.05$<br>$p = 0.852$<br>$BF_{01} = 9.38$ | $r = -0.09$<br>$p = 0.987$<br>$BF_{01} = 6.87$ | $r = -0.06$<br>$p = 0.927$<br>$BF_{01} = 8.33$ | $r = -0.33$<br>$p = 0.513$<br>$BF_{01} = 0.30$ | $r = 0.14$<br>$p = 0.784$<br>$BF_{01} = 5.61$ | 1 | |
| Pain detection | | $r = 0.04$<br>$p = 0.818$<br>$BF_{01} = 9.49$ | $r = 0.06$<br>$p = 0.945$<br>$BF_{01} = 10.02$ | $r = -0.09$<br>$p = 0.940$<br>$BF_{01} = 8.04$ | $r = -0.01$<br>$p = 0.942$<br>$BF_{01} = 6.07$ | $r = 0.12$<br>$p = 0.863$<br>$BF_{01} = 6.68$ | $r = -0.06$<br>$p = 0.980$<br>$BF_{01} = 8.92$ | 1 |

**Table 5.** Correlational matrix describing the relationship between the performances at the different interoceptive tasks (i.e., interoceptive accuracy) on the palm. Thermal interoceptive accuracy when the temperature is decreasing (cooling) is negatively correlated with cold detection, and the performance in the warm detection task is significantly correlated with both cold detection and pain detection tasks in the palm only. *P* values correspond to original FDR corrected values. *indicates *p* values that are significant after correction for multiple comparisons (FDR).

| Palm |  | Heartbeat counting task | Thermal matching task |  | Affective touch task | Temperature detection |  | Pain detection |
| --- | --- | --- | --- | --- | --- | --- | --- | --- |
|  |  |  | Increasing | Decreasing |  | Warm | Cold |  |
| Heartbeat counting task |  | 1 |  |  |  |  |  |  |
| Thermal matching task | Increasing | $r = 0.03$<br>$p = 0.851$<br>$BF_{01} = 9.67$ | 1 | | | | | |
| | Decreasing | $r = 0.16$<br>$p = 0.711$<br>$BF_{01} = 6.31$ | $r = 0.32$<br>$p = 0.105$<br>$BF_{01} = 1.2$ | 1 | | | | |
| Affective touch task | | $r = -0.04$<br>$p = 0.840$<br>$BF_{01} = 7.64$ | $r = -0.05$<br>$p = 0.865$<br>$BF_{01} = 9.55$ | $r = -0.16$<br>$p = 0.801$<br>$BF_{01} = 5.34$ | 1 | | | |
| Temperature detection | Warm | $r = 0.05$<br>$p = 0.952$<br>$BF_{01} = 9.20$ | $r = -0.20$<br>$p = 0.513$<br>$BF_{01} = 1.89$ | $r = -0.05$<br>$p = 0.921$<br>$BF_{01} = 6.29$ | $r = -0.07$<br>$p = 0.990$<br>$BF_{01} = 8.11$ | 1 | | |
| | Cold | $r = 0.01$<br>$p = 0.990$<br>$BF_{01} = 10.02$ | $r = -0.07$<br>$p = 0.990$<br>$BF_{01} = 9.15$ | $r = -0.29$<br>$p = 0.140$<br>$BF_{01} = 0.30$ | $r = -0.21$<br>$p = 0.577$<br>$BF_{01} = 2.92$ | $r = 0.41$<br>$p = 0.021^*$<br>$BF_{01} = 0.04$ | 1 | |
| Pain detection | | $r = 0.10$<br>$p = 0.950$<br>$BF_{01} = 7.31$ | $r = -0.17$<br>$p = 0.798$<br>$BF_{01} = 5.21$ | $r = -0.16$<br>$p = 0.770$<br>$BF_{01} = 6.41$ | $r = -0.33$<br>$p = 0.840$<br>$BF_{01} = 8.54$ | $r = 0.34$<br>$p = 0.042^*$<br>$BF_{01} = 0.25$ | $r = 0.12$<br>$p = 0.914$<br>$BF_{01} = 6.71$ | 1 |

The rest of the correlational analysis is reported in Table 3 for the forearm and Table 4 for the palm and is visually reported in Figure 4 in the Supplementary Materials. Performances on the thermal matching task in increasing and decreasing scales were significantly correlated only on the forearm (Table 4, Benjamini-Hochberg adjusted *p* value = 0.02), whereas the performance on the thermal matching task when temperature was decreasing was negatively correlated with cold temperature detection on the palm (Table 4, Benjamini-Hochberg adjusted *p* value = 0.14). Warm temperature detection was significantly correlated with both cold detection (Benjamini-Hochberg adjusted *p* value = 0.04) and pain (Benjamini-Hochberg adjusted *p* value = 0.01) detection in the palm, suggesting a relationship between the perception of temperature and pain detection in the palm but not in the forearm.

### Confidence across modalities

A similar approach of analysis was adopted for the confidence scores for cardiac awareness, dynamic temperature and affective touch, whereby we focused first on the different tasks separately and then on the relationship of the participants’ confidence in completing the different tasks. Confidence in one’s own performance in the *heartbeat counting task* was not related to actual performance (*r* = 0.128; *p* = 0.323, see Figure 5 in the Supplementary Materials).

In the *thermal matching task*, the results of a 2 (location) x 2 (order) repeated measures ANOVA showed a significant main effect of location (*F*(1, 61) = 24.64; *p* < 0.001), with participants being more confident with their answers in the forearm (M= 6.73; SE = 0.18) than in the palm (M = 6.22; SE = 0.20). The order of presentation of temperature (staircase increasing/decreasing) did not have a significant effect (*F*(1, 61) = 0.184; *p* = 0.67) on confidence; the interaction between staircase and location was not significant (*F*(1, 61) = 1.54; *p* = 0.22). Regarding the relationship between performance and confidence in the thermal matching task, the only significant relationship was between confidence and performance in decreasing (cooling) temperature in the forearm (*r* = 0.287; *p* = 0.025, Figure 6). All the other relationships between confidence and accuracy in the thermal matching task were non-significant (increasing forearm: *r* = 0.21; *p* = 0.10; decreasing palm: *r* = 0.19; *p* = 0.12; increasing palm: *r* = 0.04; *p* = 0.73).

**Figure 6:**
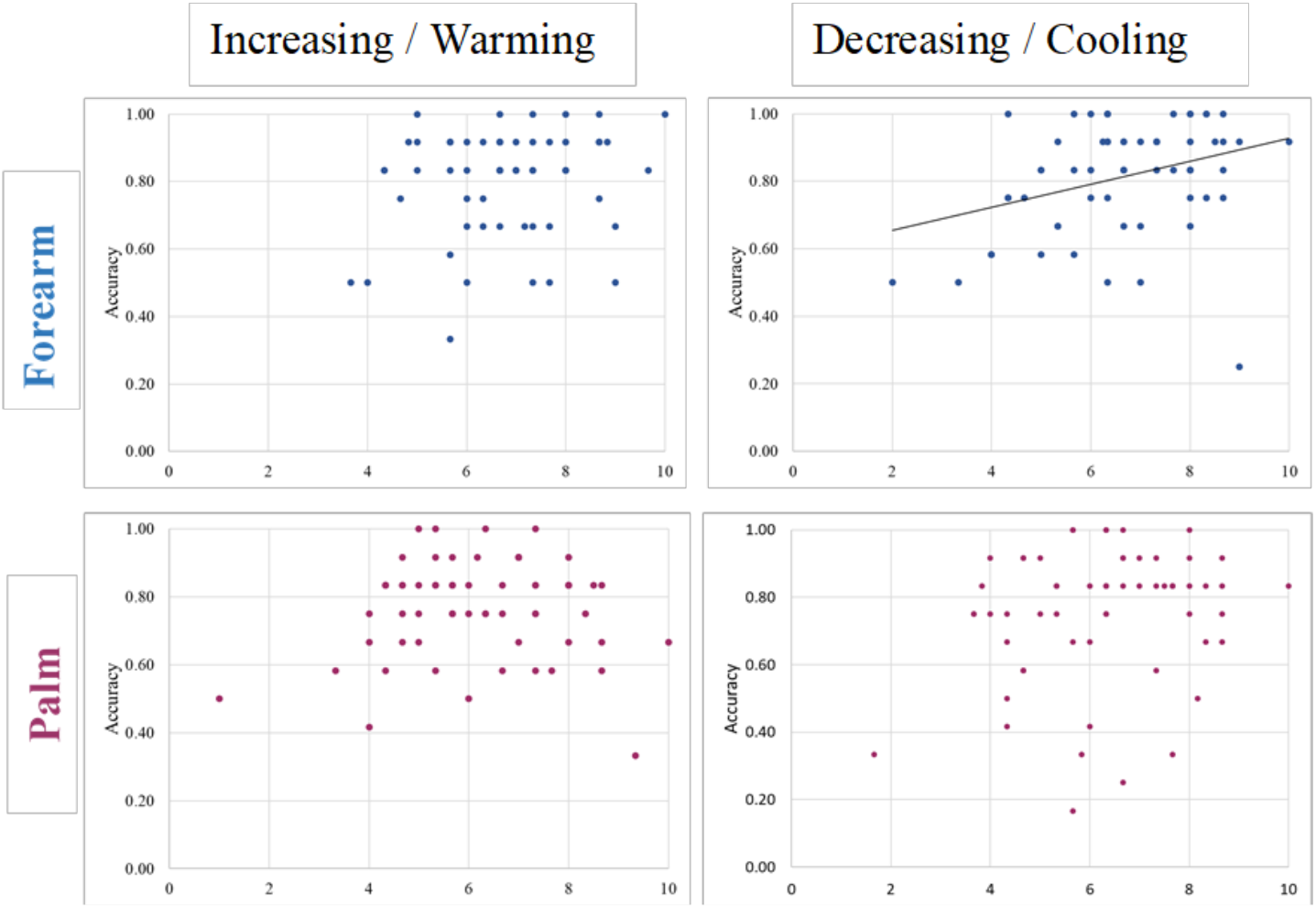
Confidence-accuracy correspondence in the thermal matching task. Only for the thermal matching task (TMT) on the forearm at decreasing (cooling) temperatures was there a correspondence between accuracy and the participants’ average confidence rating; this indicated that, at the broad group level, subjective and objective dimensions were aligned. By contrast, there was no significant relationship between confidence and accuracy for performance on the TMT regarding the palm (in burgundy).

Finally, we focused on the *affective touch task*. The results of the 2 (location) x 3 (velocities) repeated measures ANOVA showed no main effect of location (*F* (1, 61) = 0.0; *p* = 1.00) or velocity (*F* (2, 122) = 1.72; *p* = 0.183) on participants’ confidence in performance. The interaction between location and velocity was non-significant (*F* (2, 122) = 2.62, *p* = 0.077). Regarding the relationship between performance and confidence in the affective touch task, we found a significant relationship between confidence and perception of CT-optimal touch (*r* = 0.366; *p* = 0.003; Benjamini-Hochberg adjusted *p* value = 0.009) and borderline touch (*r* = 0.289; *p* = 0.023; Benjamini-Hochberg adjusted *p* value = 0.03) for the forearm only (see Figure 6 of Supplementary Materials).

Next, we investigated whether the tendency to be confident in one’s own performance accuracy was generally related across the modalities. Confidence in cardiac interoception was significantly related to confidence in thermal matching task performance for both the forearm (increasing: *r* = 0.542; *p* < 0.001; Benjamini-Hochberg adjusted *p* value = 0.01; decreasing: *r* = 0.607; *p* < 0.001; Benjamini-Hochberg adjusted *p* value = 0.01) and palm (increasing: *r* = 0.517; *p* < 0.001; Benjamini-Hochberg adjusted *p* value = 0.01; decreasing: *r* = 0.569; *p* < 0.001; Benjamini-Hochberg adjusted *p* value = 0.01). The correlations between confidence in cardiac interoception and affective touch showed a significant relationship between the former and confidence in the perception of CT-optimal touch (*r* = 0.28; *p* = 0.026; Benjamini-Hochberg adjusted *p* value = 0.035), borderline touch (*r* = 0.27; *p* = 0.034; Benjamini-Hochberg adjusted *p* value = 0.04) and CT-non-optimal touch (*r* = 0.309; *p* = 0.014; Benjamini-Hochberg adjusted *p* value = 0.02) in the palm but not in the forearm (CT optimal touch: *r* = 0.147; *p* = 0.253; borderline touch: *r* = 0.199; *p* = 0.121; CT-non-optimal touch: *r* = 0.212; *p* = 0.099).

In terms of confidence across the thermal matching task and affective touch, correlational analyses revealed a significant relationship between confidence when temperature was increasing and CT-optimal touch (*r* = 0.433; *p* < 0.001; Benjamini-Hochberg adjusted *p* value = 0.01), borderline touch (*r*= 0.448, *p* < 0.001; Benjamini-Hochberg adjusted *p* value = 0.01) and CT-non optimal touch (*r* = 0.450, *p* < 0.001; Benjamini-Hochberg adjusted *p* value = 0.01) in the palm. The same applies for confidence when the temperature was decreasing (CT-optimal touch: *r* = 0.331; *p* = 0.005; Benjamini-Hochberg adjusted *p* value = 0.01; borderline touch: *r* = 0.355; *p* = 0.005; Benjamini-Hochberg adjusted *p* value = 0.01; CT-non-optimal touch: *r* = 0.353, *p* = 0.005; Benjamini-Hochberg adjusted *p* value = 0.01) in the palm. Similar results were found for the forearm: increasing temperature and CT-optimal touch (*r* = 0.260; *p* < 0.041; Benjamini-Hochberg adjusted *p* value = 0.05), borderline touch (*r* = 0.288, *p* = 0.023; Benjamini-Hochberg adjusted *p* value = 0.03), CT-non-optimal touch (*r* = 0.328, *p* = 0.009; Benjamini-Hochberg adjusted *p* value = 0.02), and decreasing temperature and CT-optimal touch (*r* = 0.315; *p* = 0.013; Benjamini-Hochberg adjusted *p* value = 0.02), borderline touch (*r* = 0.352, *p* = 0.005; Benjamini-Hochberg adjusted *p* value = 0.01) and CT-non-optimal touch (*r* = 0.406, *p* = 0.001; Benjamini-Hochberg adjusted *p* value = 0.01).

## Discussion

### Summary of key findings

The main aims of this study were to investigate 1) thermosensation as skin-based interoception by using a novel dynamic matching task and a classic static detection task and 2) the relationships across four different interoceptive submodalities: cardiac awareness, affective touch, nociception and thermosensation. Our results showed a significantly more accurate dynamic thermal accuracy in the forearm (hairy skin) than in the palm (non-hairy skin). Moreover, the static thermal task, showed significantly lower detection thresholds for cold stimuli on the forearm than on the palm, which could indicate basic differences in cold perception between hairy and non-hairy skin. Furthermore, and in line with previous studies, participants showed a higher affective touch sensitivity, i.e., perception of the difference in pleasantness between CT-optimal and CT-non-optimal touch in the forearm compared to the palm. Interestingly, we found a significant relationship between performance on the thermal matching task (cooling) and static cold detection, as well as between static warm detection and static thermal pain only on the palm (non-hairy skin). These mixed findings point towards differences in thermal perception on hairy and non-hairy skin, with hairy skin showing more accurate thermosensation than non-hairy skin (see also Filingeri, Zhang, and Arens, 2018). Notably, no other significant correlations were observed in terms of interoceptive accuracy across modalities than the abovementioned relationships. This suggests that the submodalities of cardiac awareness, affective touch, nociception, and thermosensation are processed relatively independently. Thus, in future studies, in might be better to quantify interoception using a battery of tests to capture it fully, and we suggest caution when using the perception at one single sub-modality as a proxy for interoception generally. More details about these findings are discussed below.

### Differences in the perception of temperature and affective touch on hairy and non-hairy skin

The observed differences in the performance between hairy (forearm) and non-hairy (palm) skin could be due to fundamental differences in thermosensation on hairy and non-hairy skin, with the former activating the CT tactile system to a larger extent than the latter (Vallbo, Olausson & Wessberg, 1999; Watkins et al., 2020). These results are in line with recent findings showing higher thermal sensitivity in hairy skin than in glabrous skin (Filingeri et al., 2018). Our findings provide support for the idea that the perception of temperature on hairy skin might have a functional relationship to, at least in part, the perception of homeostatic internal signals from our body mediated by C-fibres such as classic cold and warm thermal afferents, and perhaps even CT afferents that we will discuss further below (Craig, 2009; Björnsdotter, Morrison & Olausson, 2010). If this is the case, it could be hypothesised that thermal signals detected through the hairy skin of our body might have a privileged role in social thermoregulation, maintenance of homeostasis and ultimately survival (Morrison, 2016; Burleson & Quigley, 2021), suggesting that the CT afferents might work in concert with cold and warm receptors to perceive and signal deviations from their optimal firing temperature (i.e., 32°C). Note however, that the performance on the thermal matching task and in the affective touch task was not correlated in the forearm or in the palm, which is consistent with independent processing of these signals rather than highly integrated. In contrast, we theorise that thermal signals detected through the non-hairy skin of our body (e.g., palm) might potentially have a more discriminatory role and they might therefore be important for experiencing the temperature of grasped objects, for instance, a role that is less related to thermoregulation and more to exploring the properties of external objects (Vallbo and Johansson, 1984; Johansson and Flanagan, 2009; Corniani and Saal, 2020). This is further supported by our findings showing that overall, improved performance in temperature perception on hairy skin was more consistent when the thermal stimuli were dynamic compared to that in response to static stimuli. Furthermore, we can hypothesise that thermal dynamic sensations (and in particular at neutral temperatures typical of skin to skin contact) play a different role in daily social interaction as compared to static thermal sensations. The characteristics of CT optimal stimulation (i.e., slow velocity, light pressure and neutral temperature) closely resemble the ones typical of affiliative touch (McGlone et al., 2014; Burleson and Quigley, 2021), suggesting that these interoceptive signals might promote social connection that is of vital importance for our survival. However, the thermal matching task and the temperature detection task showed a similar pattern of results in regard to the perception of cold; that is, cold detection thresholds were lower for hairy skin than for non-hairy skin when touch was both dynamic and static. The fact that we have a higher sensitivity to cooling than warming could be justified by the greater abundance of cold receptors throughout our entire body (1.3–1.6 times stronger sensitivity to cooling than to warming, Luo et al., 2020). Given the significant correlations between performance on the thermal matching task and static temperature and between performance on the thermal matching task and pain detection tasks on the palm, one potential interpretation could be the existence of overlapping processes for the perception of dynamic and static thermal signals and pain in the palm but not in the forearm, where the processing of dynamic and static thermal signals remains more segregated. This could be due to the higher density of cold and warm thermosensory fibres in hairy vs. non-hairy skin given its physiological composition (Filingeri, Zhang and Arens, 2018; Corniani and Saal, 2020) and thus – potentially – higher peripheral innervation for the detection of temperature and pain in hairy skin compared to non-hairy skin (Filingeri, Zhang and Arens, 2018).

One could argue that the observed differences between the palm and forearm might simply relate to different densities of classical cold and warm receptors in different parts of the skin. Our results from the static thermal task showing a significantly lower ability to detect cold stimuli on the forearm compared to on the palm would be consistent with this view. However, the palm of the hands and the pulps of the digits are usually engaged when performing sophisticated tasks such as object manipulation, and therefore, these areas are highly innervated with sensory afferents (Johansson and Flanagan, 2009; Vallbo and Johansson, 1984), as well as presenting differences in perception due to skin thickness, for example (Stevens & Choo, 1998). Thus, an alternative explanation and hypothesis for future studies could be that we have learnt to combine or pair painful and thermal sensations from the palm and digit when manipulating and grasping objects, objects that sometimes can cause pain. Such functional correlations can be less pronounced on the hairy skin since we do not use this skin site to actively explore objects via touch. Furthermore, we know that despite differences in the quality and quantity of nociceptors, hairy and glabrous skin share similar nociceptive afferents, with non-significant differences in pain sensations or brain potentials in terms of latency and amplitude (Iannetti, Zambreanu and Tracey, 2006). A recent study reported similar thresholds for cooling and warming in the forearm and the palm of the hand (Luo et al., 2020); however, the dorsum of the hand (hairy skin) was reported to be more thermally sensitive than the palm (non-hairy). To the best of our knowledge, it is not known whether the observed differences in sensitivity are due to the presence of fewer thermal receptors on the palm than on the forearm (but see Watkins et al., 2020 for a recent work on different CT densities in hairy and non-hairy skin). Given the privileged role of hairy skin in homeostatic maintenance of thermoneutrality and social thermoregulation (Morrison, 2016), it could be plausible that such differences in thermosensitivity correspond to a difference in the distribution of cold and warm receptors between hairy and non-hairy skin.

Ackerley et al. (2014) investigated the CT responses to tactile stimuli delivered at different speeds and temperatures; the study showed that the maximum firing rate of the CT system was recorded for mechanothermal stimuli delivered in the range of velocities between 1-10 cm/s and at neutral temperature (32°C, typical of human skin, Arens & Zhang, 2006). Thus, on the basis of this evidence and our findings, we can hypothesise that the CT system does not have an active role in signalling specific velocities and/or temperature, but it rather provides information about the affective status of the skin. This affective status is positive during affective touch and pleasant temperature. In interoceptive terms, the CT system might communicate how the skin – as an own bodily organ – affectively ‘feels’ at any given time, which is district from discriminating the somatosensory properties of external object. The fact that we did not find a relationship between performance on the thermal matching task and the affective touch task might be partially explained by recent evidence showing independent spinothalamic pathways signalling pain, temperature and itch on one hand and affective touch on the other (Marshall et al., 2019). Our additional control analysis evaluating performance on the thermal matching task in the three different temperatures 30°C, 32°C and 34°C (see Supplementary Materials) did not show any significant differences between temperatures. However, these temperatures were all in the range of thermoneutrality; thus, they might not have been sufficient to capture such differences in CT thermal specificity, suggesting that task performance might have been mainly driven by activity from cold and warm receptors. An alternative explanation could be that CT activation only reflects pleasantness and the affective aspects of touch. If this is the case, the CT contribution to performance on the thermal matching task would be neglectable; therefore, this task mainly provides information about thermosensation, regardless of the involvement of the CT system. Nevertheless, future studies could broaden the range of tested temperatures to specifically investigate the optimal thermal range of CT activation and to further clarify whether the thermal specificity of the CT afferent system relates to tactile pleasantness (thus suggesting the existence of overlapping processes), rather than the perception of temperature *per se*.

### Interoceptive accuracy across modalities

In terms of modalities, previous studies have attempted to investigate interoception across different senses and channels mainly by focusing on the comparison of such alternative modalities (e.g., gastric and respiratory functions) with performance in classic heartbeat detection tasks (e.g., Whitehead & Drescher, 1980; Herbert et al., 2012; Azzalini, Rebollo & Tallon-Baudry, 2019; Garfinkel et al., 2016; Ferentzi et al., 2018; Faull, Subramanian, Ezra & Pattinson, 2019; Monti, Porciello, Tieri & Aglioti, 2020). The performance on such tasks seems to be dissociated from cardiac accuracy. A few studies also attempted to investigate the relationship between skin-mediated interoceptive signals and cardiac awareness, showing no significant relationship between pain perception (Werner et al., 2009; Ferentzi et al., 2018) and affective touch (Crucianelli et al., 2018) with respect to performance on the heartbeat counting task.

In keeping with these studies, our results confirm and further extend this lack of relationship between skin-mediated interoceptive modalities and cardiac awareness. This suggests that cardiac interoception, pleasant touch, thermosensation and nociception are working to a large extent as independent sensory modalities, which contrasts with the view of interoception as a single or highly integrated perceptual function. Thus, our main findings regarding the actual performance on interoceptive tasks (i.e., accuracy) are in keeping with previous studies highlighting that interoceptive accuracy assessed via a single modality cannot be generalised across other channels (e.g., no relationship between cardiac interoception, gastric perception, pain, taste, Ferentzi et al., 2018; no relationship between cardiac interoception and respiration, Garfinkel et al., 2016; and no relationship between cardiac interoception and affective touch, Crucianelli et al., 2018, but see Herbert et al., 2012 for a negative correlation between cardiac interoception and gastric function). Here, we investigated for the first time the relationship between multiple externally generated, skin-mediated interoceptive modalities, including thermosensation. Our findings on interoceptive accuracy combined with the aforementioned ones provide stronger and more conclusive behavioural evidence of the independence of different interoceptive sub-modalities. In particular, this study shows for the first time that this is the case even when the interoceptive modalities are mediated by the same source, that is, the skin. Thus, it could be argued that the best way to characterise individual interoceptive abilities is by means of an ‘interoceptive battery’, which includes and targets a variety of modalities, rather than using one single task and then generalising the results to the entire construct of interoception. Although the central processing of interoception signals is associated with the posterior and anterior insula in the literature (e.g., pain, Segerdahl et al., 2015; temperature, Craig et al., 2000, and pleasant touch, Björnsdotter et al., 2011), parallel and separate pathways are involved in the peripheral processing of such signals (e.g., Marshall et al., 2019), and it is not surprising that such processes can remain mainly independent.

### Interoceptive ability across dimensions

In terms of the *metacognitive dimension*, our results show a dissociation between beliefs of performance (metacognitive awareness) and actual performance across the different interoceptive modalities, in line with previous studies that failed to find a relationship between these two dimensions of interoception (e.g., Garfinkel et al., 2016), with the exception of cardiac interoceptive accuracy, for which the prior belief of performance was a significant predictor (as in Ring & Brener, 1996). Thus, a multidimensional assessment of interoception that characterises interoception both at the level of objective perception and at the level of metacognition might be necessary to gain a more complete picture both at the perceptual level and the metacognitive level (Garfinkel et al., 2015; Quadt, Critchley & Garfinkel, 2019).

In keeping with previous studies, our results highlight a general relationship between the beliefs regarding performance across tasks (as in Beck et al., 2019); that is, people who had higher beliefs of performance in the heartbeat counting task or thermal matching task also had higher beliefs of performance in the affective touch and pain detection task. This might reflect the fact that this dimension of interoception is related to domain general cognitive abilities and therefore is mainly driven by top-down beliefs that are independent from perceptual interoception performance.

Similarly, the confidence in performance was correlated across modalities; in particular, the tendency to be confident in one’s own performance showed significant linear relationships between the thermal matching task and both cardiac interoception and affective touch. Higher confidence was also related to better performance in both the thermal matching task and the affective touch task in hairy skin (forearm) only, in keeping with recent findings arguing that such ‘metacognitive sensitivity’ is higher in hairy skin than in non-hairy skin (von Mohr, Kirsch, Loh & Fotopoulou, 2019). Overall, participants performed significantly better on the thermal matching task when hairy skin was involved than when non-hairy skin was involved, and this enhanced performance was related to a higher level of confidence in their performance. This evidence might suggest that we are more precise in or aware of our ability to detect such stimuli on hairy skin for affiliative and thermoregulatory reasons (Morrison, 2016; Filingeri, Zhang & Arens, 2018), processes that could have been linked to the existence of fur on the body in the past. Taken together and in contrast with the interoceptive accuracy results reported above, the results of the confidence levels across interoceptive modalities suggest the existence of a common ability or system that we use when judging our interoceptive abilities at a metacognitive level. In other words, confidence ratings share common metacognitive task demands that are not modality specific but are rather generalisable.

In terms of *interoceptive sensibility*, here we used both the Body Awareness Questionnaire (Shields, Mallory & Simon, 1989) and the Body Perception Questionnaire (Porges,1993). Our results showed a correlation between the responses to the two questionnaires; however, these seem to be distinct in the way they relate to interoception. In particular, the Body Awareness Questionnaire seems to relate more to the metacognitive dimension of interoception (both in terms of prior beliefs of performance and confidence, see Table 2 in Supplementary Materials) rather than to objective performance on interoceptive tasks (i.e., accuracy), for which no significant correlations were found. In contrast, the Body Perception Questionnaire was significantly correlated with actual performance in the affective touch task on the palm only (see Table 3 in Supplementary Materials). The Body Perception Questionnaire was also positively correlated with confidence in the affective touch task in both the palm and forearm, as well as prior belief in performance in pain detection only, suggesting a relationship between this self-report measure and metacognitive aspects of interoception (see Garfinkel et al., 2015; Fairclough & Goodwin, 2007; Schulz et al., 2013 for similar approaches). We believe that these findings might inform future studies that aim to include a measure of interoceptive sensibility. For example, it could be valuable to use the recently developed and standardised Multidimensional Assessment of Interoceptive Awareness scale (Mehling, Price, Daubenmier, Acree, Bartmess & Stewart, 2012), which is organised in eight separate subscales (e.g., emotional awareness, body listening, and self-regulation) and could potentially target different facets of self-reported interoception and their possible relationships with interoceptive accuracy and metacognitive awareness. Another valuable alternative is the newly developed Interoceptive Accuracy Scale (IAS, Murphy et al., 2020), which aims to distinguish the attention component from the accuracy one in interoceptive self-report measures.

Even though we did not find significant relationships between the questionnaires and the interoceptive accuracy measures in the present study, this does not mean that the former does not provide important information or should not be used in future studies. For instance, interoceptive sensibility has proven to be clinically relevant since some individuals have shown a dissociation between their self-report abilities to perceive their body and actual performance, such as in individuals with autism spectrum disorder, individuals with high levels of anxiety (e.g., Garfinkel et al., 2016) and individuals with an eating disorder (Pollatos and Georgiou, 2016; Eshkevari et al., 2014).

### Limitations and future directions

Investigations around thermosensation as an interoceptive modality have been sparse. One reason could lie in the difficulties encountered when focusing on the perception of temperature, mainly because tactile and temperature stimuli often occur simultaneously in everyday life, with temperature being an intrinsic property of touch carrying both interoceptive and exteroceptive information (i.e., every tactile stimulation provides thermal sensation, and the source of the tactile stimulation can be cooler, the same or warmer than the skin temperature). Thus, future studies could attempt to investigate the interoceptive nature of the perception of temperature over and above the perception of touch to try to disentangle thermosensation from tactile somatosensation. For example, stimulating thermosensors by means of heat lamps or lasers can allow us to deliver contactless thermal signals, and address questions such as “which limb feels warmer” rather than discriminating the temperature of an object touching the skin. Nevertheless, we strongly believe that the present work provides an important first step towards a better understanding of the interoceptive nature of some thermosensory tactile stimuli, which combined with the results of a contactless approach, would provide complementary knowledge necessary to gain a better understanding of thermal interoception.

To our knowledge, this is the first experimental study to investigate the perception of dynamic thermal stimuli with a particular focus on hairy and non-hairy skin. Therefore, future studies should focus on validating the thermal matching task by exploring more body sites, adding more trials and investigating the test-retest reliability of the task. In the thermal matching task, we used a 3-second interval between trials, and it could be argued that this interval might not be sufficient for the skin to return to its starting temperature. We did not record the skin temperature continuously during the task, but only at the beginning, and we acknowledge this as a potential limitation of the study. However, our data showed no significant variations between skin sites and across participants at baseline, so we believe that this factor unlikely played a crucial role.

Our study investigated individual differences in four different interoceptive submodalities, and suggested that these seem to be relatively independent. However, we have not targeted the mechanisms behind such interindividual variability in interoception. One could speculate that individual differences in interoceptive sub-modalities could be driven by peripheral factors, such as different receptor densities, or by central differences, such as higher cognitive and top-down factors, central pathways, grey matter thickness in cortical areas and decision processes. Future studies could explore the underlying mechanisms of such effects further and provide more insight into this matter. For example, future ad hoc physiological studies could investigate the performance at the thermal matching task using a static stimulation instead or focus on the perception of temperature in relation to the density of CT and non-CT thermal receptors in hairy and non-hairy skin. In this context, it would also be interesting to explore the relationship between the thermal matching task and cold pain sensitivity, for instance, since here we only investigated heat pain. It might also be interesting to provide more insight into the integration of different interoceptive modalities at the central level to better specify the interoceptive pathways of the different interoceptive sub-modalities. For example, it would be interesting to also consider tactile pleasantness in relation to the stimulation provided as part of the thermal matching task. Being that temperature perception is mediated by the skin and is therefore occurring at the border between inside and outside the body, future studies should focus on the perception of dynamic temperature in relation to classic models of multisensory integration and body representation. This approach would provide important insight into the way the brain weighs interoceptive and exteroceptive signals to build a unitary and coherent sense of self.

## Conclusions

Taken together, our results suggest that it is possible to potentially broaden the testable interoceptive modalities beyond cardiac signals to include temperature perception, as tested by means of our newly developed thermal matching task. The perception of dynamic temperature can provide information about our ability to detect externally generated skin-mediated signals that translate into internal, physiological sensation. The accuracy in detecting such stimuli might relate to the ability to react promptly to potential threats to our thermoneutrality, thus reflecting the way we perceive and respond to homeostatic needs. This study provides further support to the existence of different processes underlying the perception of temperature on hairy and non-hairy skin, with hairy skin having higher thermosensory peripheral innervation and potentially playing a more critical role in interoceptive aspects of thermosensation as well as thermoregulation (see also Filingeri, Zhang & Arens, 2018). The lack of significant relationships between performance on interoceptive tasks across all the modalities tested (i.e., cardiac accuracy, affective touch, temperature and pain detection), supports the idea that interoception might be better conceptualised as a modular construct with relatively independent processing in parallel streams. Thus, just as in the case of exteroception, distinct interoceptive modalities might not necessarily be related to one another. Consequently, future basic and clinical studies could benefit from using interoceptive batteries of tests that comprise multiple interoceptive modalities, including the thermal matching task, which could collectively provide a complete overview of interception in health and disease.

## Acknowledgements

The authors would like to thank Martti Mercurio and Bo Johansson for their assistance with the equipment, and Dr Rochelle Ackerley for useful insight in data interpretation. This work was supported by the Göran Gustafsson foundation, the Swedish Research Council (Distinct Professor Grant) and the European Research Council (SELF-UNITY). Laura Crucianelli was supported by the Marie Skłodowska-Curie Individual Fellowship (Homeothermic Self).

## Authors contribution

Conceptualisation: LC, HHE; Data curation: LC, AE; Formal analysis: LC; Funding acquisition: HHE, LC; Investigation: LC, AE; Methodology: LC, HHE; Project administration: LC, HHE; Resources: HHE; Visualisation: LC, AE; Writing – original draft: LC; Writing – review & editing: LC, AE, HHE

