## Supplementary Materials for "Probing interoception via thermosensation: No specific relationships across multiple interoceptive sub-modalities"

### Skin temperature values

**Table 1**. Skin temperature values recorded at the beginning of the experimental tasks. Location on the body goes from proximal (1) to distal (3). In the case of the outer forearm we started from proximal to the elbow (1), half way between elbow and wrist (2) and wrist (3). In the case of the palm, we started from proximal to the wrist (1), middle palm (2) and the base of the fingers (3). Values reported are mean and standard deviations in parenthesis.

| | **THERMAL MATCHING TASK** | | **TEMPERATURE DETECTION TASK** | |
|---|---|---|---|---|
| Location | Forearm | Palm | Forearm | Palm |
| 1 | 35.67 (0.55) | 35.63 (0.68) | 35.23 (0.51) | 35.66 (0.64) |
| 2 | 35.56 (0.56) | 36.25 (1.62) | 35.28 (0.53) | 36.00 (0.79) |
| 3 | 35.42 (0.72) | 35.89 (0.85) | 35.24 (0.75) | 35.91 (0.76) |

#### Results

#### 1. Thermal Matching Task: Control analysis

Given the evidence showing optimal CT activation at 32°C (Ackerely et al., 2014), we conducted a control analysis to explore whether there was a difference in the performance at the thermal matching task on hairy and non-hairy skin based on the reference temperature used each trial (30, 32 and 34°C). The results of the 3 (temperature) x 2 (scale: increasing vs. decreasing) x 2 (skin site: hairy vs. non-hairy) revealed a significant main effect of skin location ($F$ (1, 60) = 6.52, $p$ = 0.01), with higher accuracy in performance in hairy (M of error = 1.48; SE = 0.129) compared to non-hairy skin (M of error = 1.86; SD = 0.134). There was no main effect of scale ($F$ (1, 60) = 0.09, $p$ = 0.77) nor of temperature ($F$ (2, 120) = 0.19, $p$ = 0.82). However, there was a significant interaction between temperature and scale ($F$ (2, 120) = 4.89, $p < 0.01$). Bonferroni-corrected post hoc analysis showed a significant difference only between the performances at the task at 30°C and 34°C when temperature was decreasing ($t$ (61) = -2.60, $p$ = 0.01). Finally, there was no significant interaction between temperature and location ($F$ (2, 120) = 0.90, $p$ = 0.41) nor between scale and location ($F$ (1, 60) = 0.67, $p$ = 0.42). The three-way interaction between temperature, scale and location was non-significant ($F$ (1, 120) = 0.13, $p$ = 0.88). It should be noted that the present thermal matching task used temperatures in the range of neutral temperature; thus, they might have not been sufficient to highlight differences in CT thermal sensitivity. Future studies could broad the range of tested temperature to specifically investigate the optimal thermal range of CT activation.

### 2. Affective touch task

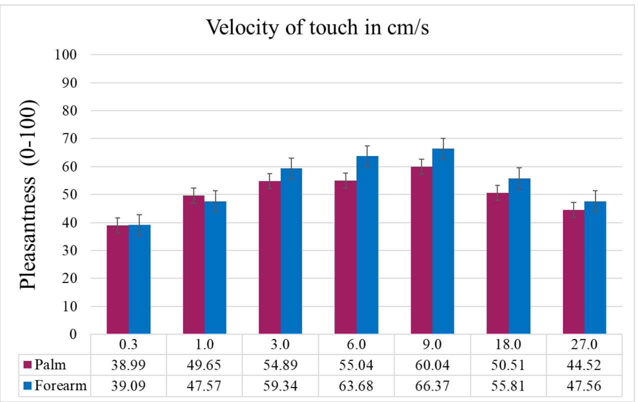


**Figure 1.** Mean and standard errors for the affective touch task for each velocity (velocities are reported in cm/s), showing the main effect of velocity and location (forearm vs. palm) on touch pleasantness. The average ratings of perceived pleasantness in response to brush stroking are in line with the ones reported by Löken et al., 2009.

### 3. (Static) pain detection task

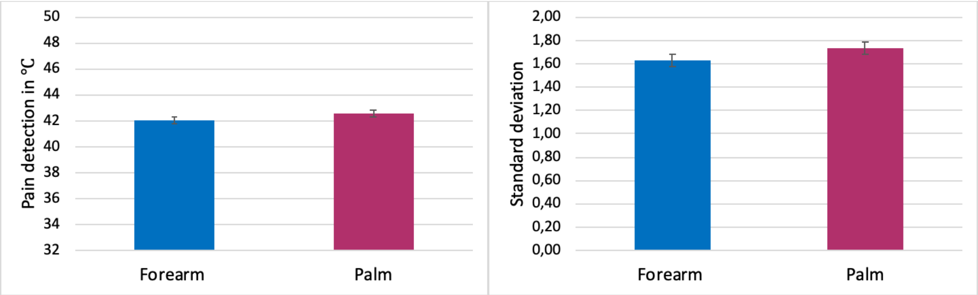


**Figure 2**. Mean and standard errors for the static pain detection task, showing no significant difference between palm and forearm in the detection of pain (quantified as temperature) and consistency in detecting pain (quantified as standard deviation).

### 4. Interoceptive accuracy across modalities

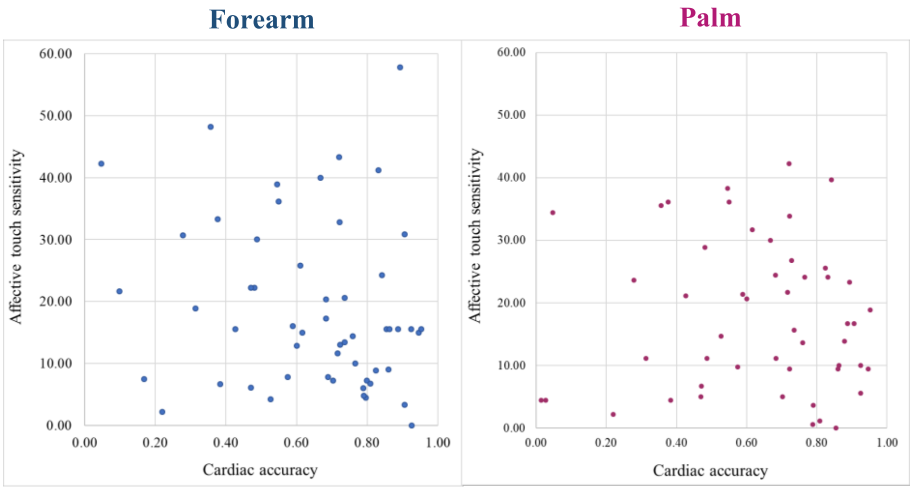


**Figure 3.** Scatter plot representing the non-significant relationship between the performance at the heartbeat counting task (cardiac accuracy) on the x-axis and the affective touch sensitivity in the forearm and palm.

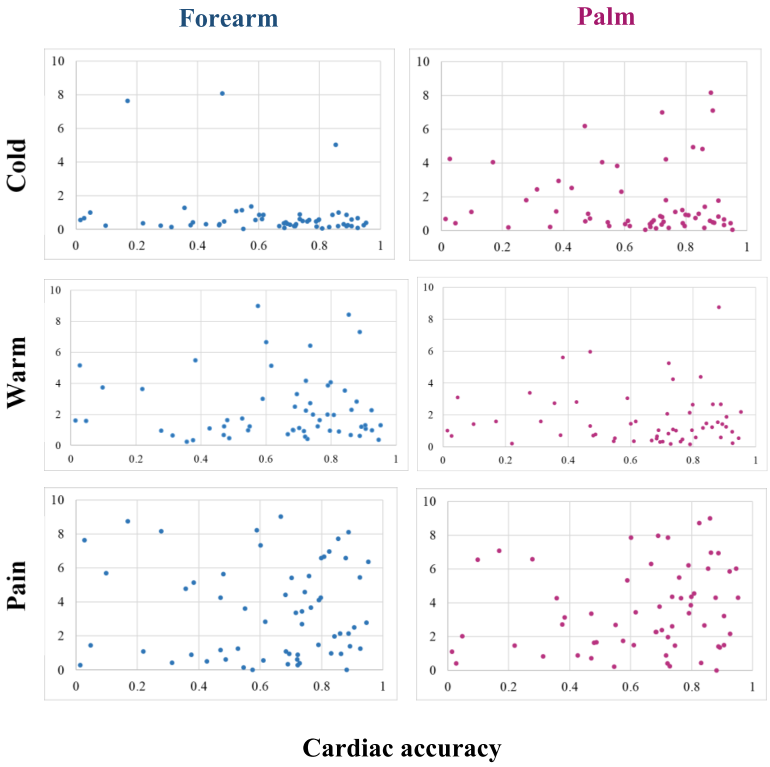


**Figure 4.** Scatter plot representing the non-significant relationship between the performance at the heartbeat counting task (cardiac accuracy) on the x-axis and the static temperature and pain in the forearm and palm.

### 5. Confidence in performance across modalities

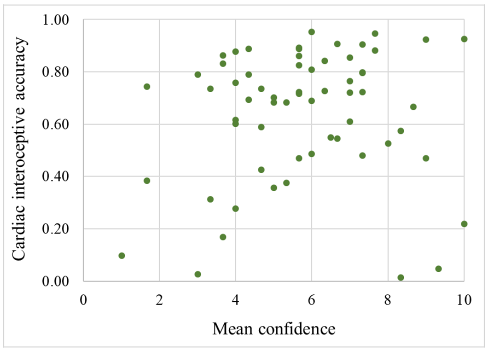


**Figure 5.** Confidence-accuracy correspondence in the Heartbeat Counting Task.

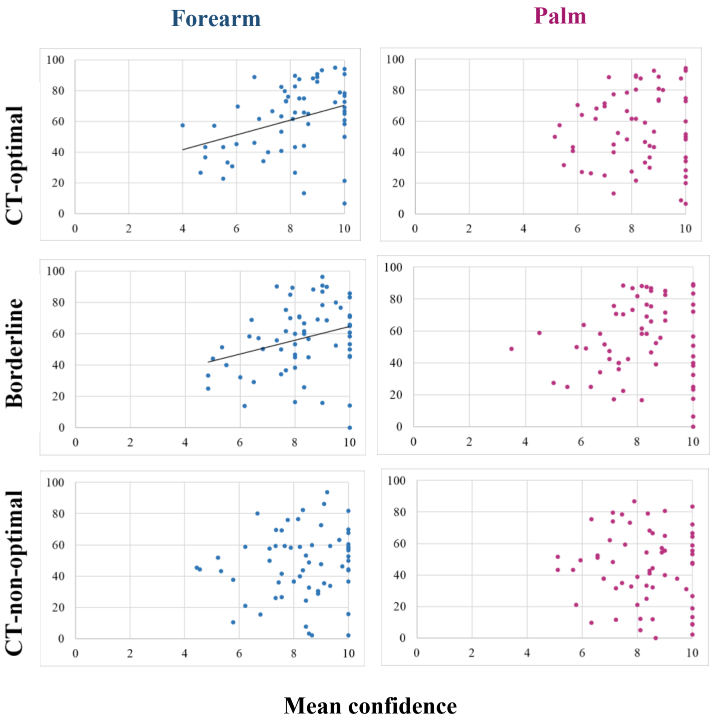


**Figure 6.** Confidence-accuracy correspondence in the Affective Touch Taks. In the y-axis the tactile pleasantness at CT-optimal, borderline and CT-non-optimal velocities are reported for both forearm and palm.

### 6. Prior metacognitive beliefs about performance across modalities

We investigated whether prior beliefs about the performance could predict the performance at the tasks. For the *cardiac interoception*, a significant regression equation was found (R = 0.35; $R^2$ = 0.123; $F$ (2, 61) = 4.13; $p$ = 0.02) with only the prior belief explaining 12.3 % of the variance in the performance at heartbeat counting task.

For the *affective touch task*, prior belief of performance was not a significant predictor of performance at the task on the forearm (R = 0.349, $R^2$ = 0.122, F (4, 61) = 1.975; p = 0.111) nor on the palm (R = 0.179, $R^2$ = 0.032, F (4, 61) = 0.472; p = 0.756).

For the *thermal matching task*, a non-significant regression equation was found for the increasing temperature in the palm (R = 0.045; $R^2$ = 0.002; $F$ (2, 61) = 0.059; $p$ = 0.943) and in the forearm (R = 0.246; $R^2$ = 0.061; $F$ (2, 61) = 1.90; $p$ = 0.158). Similarly, a non-significant regression equation was found for the increasing temperature in the palm (R = 0.045; $R^2$ = 0.002; $F$ (2, 61) = 0.059; $p$ = 0.943) and in the forearm (R = 0.246; $R^2$ = 0.061; $F$ (2, 61) = 1.90; $p$ = 0.158).

Finally, we investigated whether there was a significant relationship between the prior beliefs of performance across modalities. Results of the correlational analysis showed that the prior belief of performance at heartbeat counting task was positively correlated to the belief about pain detection ($r$ = 0.429; $p$ = 0.001) and temperature detection ($r$ = 0.523; $p$ < 0.001). Similarly, the belief of performance at the affective touch was significantly correlated with both pain ($r$ = 0.444; $p$ < 0.001) and temperature ($r$ = 0.294; $p$ = 0.020) detection. Also, the belief of performance in pain detection was related to the one for temperature detection ($r$ = 0.575; $p$ < 0.001). Finally, the belief of performance at the thermal matching task was significantly related to the one for temperature detection ($r$ = 0.37, $p$ = 0.003).

### 7. Interoceptive sensibility across modalities

The Body Awareness Questionnaire (BAQ) and the Body Perception Questionnaire (BPQ) were significantly correlated to each other ($r$ = 0.61; $p$ < 0.01).

The Body Awareness Questionnaire (BAQ) was not related to any of the *interoceptive accuracy* (i.e. actual performance at the task) across modalities (see Table 2). However, the BAQ was related to the *confidence* about performance in the cardiac interoception ($r$ = 0.355; $p$ = 0.005); in the thermal matching task in both palm (increasing: $r$ = 0.390; $p$ = 0.002; decreasing: $r$ = 0.419; $p$ = 0.001) and forearm (increasing: $r$ = 0.428; $p$ = 0.001; decreasing: $r$ = 0.358; $p$ = 0.004), and the affective touch task on the palm only (CT-optimal touch: $r$ = 0.251; $p$ = 0.049; CT-non-optimal touch: $r$ = 0.258; $p$ = 0.043).

**Table 2.** Correlational matrix describing the relationship between the performances at the different interoceptive tasks (i.e., interoceptive accuracy) on the forearm and palm and the Body Awareness Questionnaire (i.e., interoceptive sensibility).

| | **Heartbeat counting task** | **Body site** | **Thermal matching task** | | **Affective touch task** | **Temperature detection** | | **Pain detection** |
|---|---|---|---|---|---|---|---|---|
| | | | **Increasing** | **Decreasing** | | **Warm** | **Cold** | |
| **Body Awareness Questionnaire** | $r$ = 0.08<br>$p$ = 0.52 | *Forearm* | $r$ = 0.15<br>$p$ = 0.52 | $r$ = 0.09<br>$p$ = 0.51 | $r$ = 0.01<br>$p$ = 0.97 | $r$ = -0.01<br>$p$ = 0.96 | $r$ = 0.01<br>$p$ = 0.97 | $r$ = -0.15<br>$p$ = 0.25 |
| | | *Palm* | $r$ = -0.06<br>$p$ = 0.64 | $r$ = -0.01<br>$p$ = 0.93 | $r$ = -0.08<br>$p$ = 0.52 | $r$ = 0.12<br>$p$ = 0.35 | $r$ = 0.18<br>$p$ = 0.16 | $r$ = - 0.10<br>$p$ = 0.46 |

Finally, we investigated the relationship between BAQ and *metacognitive beliefs* about performance. Results revealed a significant relationship between BAQ and prior belief of performance in temperature detection only ($r = 0.340$; $p = 0.007$).

Similarly, the Body Perception Questionnaire (BPQ) was not related to the *interoceptive accuracy* (i.e., actual performance at the task) across modalities, beside the affective touch sensitivity on the palm (see Table 3).

**Table 3.** Correlational matrix describing the relationship between the performances at the different interoceptive tasks (i.e., interoceptive accuracy) on the forearm and palm and the Body Perception Questionnaire (i.e., interoceptive sensibility).

| | Heartbeat counting task | Body site | Thermal matching task | | Affective touch task | Temperature detection | | Pain detection |
|---|---|---|---|---|---|---|---|---|
| | | | Increasing | Decreasing | | Warm | Cold | |
| Body Perception Questionnaire | $r = 0.21$<br>$p = 0.10$ | *Forearm* | $r = 0.10$<br>$p = 0.44$ | $r = 0.13$<br>$p = 0.30$ | $r = 0.07$<br>$p = 0.56$ | $r = -0.01$<br>$p = 0.96$ | $r = -0.18$<br>$p = 0.16$ | $r = -0.16$<br>$p = 0.23$ |
| | | *Palm* | $r = 0.04$<br>$p = 0.79$ | $r = 0.08$<br>$p = 0.51$ | **$r = -0.28$<br>$p = 0.03$*** | $r = 0.11$<br>$p = 0.39$ | $r = 0..12$<br>$p = 0.34$ | $r = -0.17$<br>$p = 0.19$ |

In contrast to the BAQ, the BPQ was related to the *confidence* about performance in the affective touch in both palm (CT-optimal touch: $r = 0.368$; $p = 0.003$; borderline touch: $r = 0.362$; $p = 0.004$; CT-non-optimal touch: $r = 0.376$; $p = 0.003$) and forearm (CT-optimal touch: $r = 0.367$; $p = 0.003$; borderline touch: $r = 0.351$; $p = 0.005$; CT-non-optimal touch: $r = 0.426$; $p = 0.001$). Finally, we investigated the relationship between BPQ and *metacognitive beliefs* about performance. Results revealed a significant relationship between BAQ and prior belief of performance in pain detection only ($r = 0.324$; $p = 0.010$).
